# Beauty That Moves: Dance for Parkinson’s Effects on Affect, Self-Efficacy, Gait Symmetry and Dual Task Performance

**DOI:** 10.1101/2020.11.23.394163

**Authors:** Cecilia Fontanesi, Joseph FX DeSouza

## Abstract

**Background:** Previous studies have investigated the effects of dance interventions on Parkinson’s motor and non-motor symptoms in an effort to develop an integrated view of dance as a therapeutic intervention. This within-subject study questions whether dance can be simply considered a form of exercise by comparing a Dance for Parkinson’s class with a matched-intensity exercise session lacking dance elements like music, metaphorical language, and social reality of grace and beauty.

**Methods:** In this repeated-measure design, 7 adults with Parkinson’s were tested four times; (i) before and (ii) after a Dance for Parkinson’s class, as well as (iii) before and (iv) after a matched-intensity exercise session. Physiological measures included heart rate and electrodermal activity. Self-reported affect and body self-efficacy were collected. Gait symmetry and dual task cost were analyzed using the 6-minutes walking test (6MWT) and Timed-Up-and-Go test (TUG), respectively.

**Results:** Average heart rate was the same for both conditions, while electrodermal activity was higher during Dance for Parkinson’s. Significant differences were found in body self-efficacy, beauty subscale, symmetry of gait, and dual task performance.

**Conclusions:** Dance, compared to an exercise intervention of matched intensity, yields different outcomes through the means of intrinsic artistic elements, which may influence affective responses, the experience of beauty, self-efficacy, and gait performance.

## 1 Introduction

Parkinson’s disease (PD) is a progressive neurodegenerative condition characterized by motor and non-motor signs. The former include tremor, bradykinesia, rigidity, gait dysfunction, and postural instability, while the latter might include depression, executive dysfunction, and sleep disorders. PD has been associated with dopamine depletion in the nigrostriatal pathway, with a substantial loss of pigmented dopaminergic cells in the substantia nigra pars compacta. The specific causes and mechanisms of this relentless degeneration are not completely understood, and there is currently no cure available besides pharmacological treatment to help patients coping with symptoms (palliative care). Physical therapy and aerobic exercise interventions can be very helpful in promoting mobility, helping reducing falls, and improving gait and balance (Frazzitta et al., 2013; Petzinger et al., 2013). Since 2007, several studies investigated the effects of dance in the Parkinson’s population, as dance practice has become increasingly popular among people living with PD.

Most studies on the effects of dance in Parkinson’s have focused on changes in motor performance, showing increased balance (Hackney and Earhart, 2009; Hackney and Earhart, 2010; Duncan and Earhart, 2012; Volpe et al., 2013; Houston and McGill, 2013; Sharp and Hewitt, 2014), improved sit-to-stand and timed-up-and-go performance, endurance and walking velocity (Volpe et al., 2013; de Natale et al., 2017; Hackney and Earhart, 2009; Hackney and Earhart, 2010; Duncan and Earhart, 2012; Sharp and Hewitt, 2014; Bearss et al., 2017; de Natale et al., 2017; Kunkel et al., 2017), and reduction in freezing of gait after several weeks of dance training (Hackney et al., 2007; Heiberger et al., 2011; Volpe et al., 2013).

Also, a few researchers have emphasized the importance of investigating non-motor effects of dance practice in PD, such as cognitive, emotional, and social benefits (McGill et al., 2014; Ciantar et al., 2019) showing changes in spatial cognition (McKee and Hackney, 2013; de Natale et al. 2017), mood (Hashimoto et al., 2015; Kunkel et al., 2017), quality of life (Bearss et al. 2017; Westheimer et al., 2015), self-efficacy and identity (Koch et al., 2016), participation and social-connectedness (Foster et al., 2013; Houston and McGill, 2013; Rocha et al., 2017). The range between studies that focus on motor outcomes (Sharp and Hewitt, 2014) and non-motor outcomes (McNeely et al., 2015) may reveal an attempt to understand the multifaceted implications of dance as a therapeutic intervention for people with PD. However, in separating these, a gap is introduced in viewing dance as either primarily a form of exercise, thus the focus on motor outcomes, or a form of psychosocial support, prioritizing non-motor effects. Although some efforts have been made to examine the interplay between these levels (Houston and McGill, 2013; de Natale et al. 2017), not many studies have achieved an integrated view yet.

Importantly, in a 2-year follow-up study (Frazzitta et al., 2015), an intensive rehabilitation and exercise treatment was found to slow down the progression of motor symptoms, reducing the need for an incremental drug dosage overtime. In parallel to this study, there are preliminary indications that dance can be disease modifying over 3-years using UPDRS as a clinical marker (DeSouza and Bearss, 2018). However, a phase 2 randomized clinical trial, Schenkman et al. (2018) emphasized that high-intensity exercise (treadmill training) could modify disease severity in PD, while moderate-intensity exercise had no effect. Thus, it remains to be understood why the reported effects of dance practice in Parkinson’s include a broad range of motor benefits, from balance to gait performance and functional mobility, since it’s rare that dance interventions for people with PD reach high-intensity aerobic levels. We need to parcel out the element that make dance an effective intervention and not simply reducible to a form of physical exercise.

Recent studies comparing the effects of dance classes and exercise interventions in healthy seniors (age 63-80 year) showed structural MRI and neurotrophic factors changes, balance, attention and memory scores that favored dance over traditional health fitness training at both 6-months (Rehfeld et al., 2018) and 18-months (Rehfeld et al., 2017; Müller et al., 2017). However, the authors do not attempt to disentangle the elements that make dance more effective than a traditional exercise intervention of similar intensity (Rehfeld et al., 2018). It is necessary to question which are the specific factors that may be responsible for dance’s therapeutic role, and how these factors may be relevant to people with PD. This study explores the hypothesis that dance, compared to an exercise intervention of matched intensity, may yield different non-motor and motor outcomes because of intrinsic artistic factors, using the same patients as their own controls. Dance elements like music, metaphorical language, and a shared reality of grace and beauty, supported by dance teachers, live musicians, and peers, may be otherwise viewed as influencing the feasibility of the interventions and participants’ compliance. However, the specific relationship to movement elicited by these elements inherent in dance may be responsible for the performance improvement and the modulation of symptoms in people with PD. In particular, the study hypothesis is that Dance for PD classes through the use of music, metaphorical language, and a socially reinforced reality of grace and beauty may produce physiological, affective, self-efficacy, and motor changes that differ from a matched-intensity exercise lacking these dance elements.

Importantly, in this study we designed an ad-hoc control intervention to compare a Dance for PD class to a movement intervention matching dance in intensity and structure but lacking the aforementioned artistic elements inherent to dance, within the same subjects. Previous studies reported the effects of dance in Parkinson’s in the absence of controls (Heiberger et al., 2011; Houston and McGill, 2013; Westheimer et al., 2015; Koch et al., 2016; Bearss et al. 2017), compared to age matched controls (Ciantar et al., 2019), using different dance styles (Hackney et al., 2007; Hackney and Earhart; 2009, Hackney and Earhart, 2010; McNeely et al., 2015; Rocha et al., 2017), as well as comparing dance to no-intervention or usual care (Duncan and Earhart, 2012; Foster et al., 2013; Hashimoto et al., 2015; Kunkel et al., 2017), standard physiotherapy or rehabilitation exercises (Volpe et al., 2013; Hashimoto et al., 2015; de Natale et al. 2017), and education (McKee and Hackney, 2013). The choice of an appropriate comparison is necessary to question the specific factors that may be responsible for the therapeutic effects of dance classes. We questioned whether dance might yield different outcomes because of factors inherent in this artistic practice. Therefore, we manipulated the experimental conditions to match subjects, location, time of the day, aerobic intensity, class structure and progression, while removing dance elements like music, metaphorical language, dance teachers and dance peers supporting an experience of grace and beauty. This comparison is necessary to investigate how these factors may be relevant to people with PD, affect non-motor and motor symptoms, thus, eventually playing a role in participants’ performance.

Rehabilitation studies may investigate both acute and long-term effects of therapeutic interventions (Bryant et al., 2016) to examine how treatment improves function immediately after a session as well as after successive applications. In populations served by community-based organizations, like Dance for PD, walk-ins are common and deeper knowledge of the effectiveness of a single-session may support practitioners’ understanding (Hymmen et al., 2013). The first study investigating the short-term, acute effects of a single Dance for PD class (Heiberger et al., 2011) explored both motor performance and quality of life. The study reported a significant effect immediately after a single Dance for PD class on functional mobility, rigidity scores, hand movements, finger taps, and facial expression. Participants also reported a beneficial effect on quality of life, which was paralleled by their caregivers. Similarly, Koch et al. (2016) reported increased well-being, body self-efficacy, and experience of beauty after a single Argentine tango class for people with PD, while both balance and gait performance were found improved after a single Dance for PD session (Bearss et al., 2017). The effects of single dance classes are important to investigate in “real-world” community settings, in which participants are welcome on a walk-in basis. Further, understanding these short-term effects can inform the rationale for structuring and dosing successive applications that aim to produce longterm effects.

## 2 MATERIALS AND METHODS

### 2.1 Study Development and Ethical Approval

The present research constitutes a repeated-measure design study on the acute effects of a dance class in people with Parkinson’s Disease (PD). Four institutions participated in this study: The City College of New York (CUNY), University of Brescia (Italy), York University (Toronto, Canada), and The Mark Morris Dance Center (Brooklyn). The design and methodology of this study were approved in February 2018 by The City College of New York Institutional Review Board (IRB).

### 2.2 Recruitment

The within-subjects study design focused on active members of the Dance for Parkinson’s community at The Mark Morris Dance Center (MMDC), who regularly took part in the Dance for PD classes. The subjects’ participation in this study was voluntary, and the subjects did not receive any compensation. The inclusion criteria were an age between 55 and 85 and a diagnosis of Parkinson’s Disease (PD) or Parkinsonism. The exclusion criteria were the inability to understand and communicate in English. During the data collection period, a total of 7 subjects were recruited, stayed in the study and in the analyses. All 7 subjects completed the baseline assessment and were tested for outcome measures at four time points, (i) before and (ii) after Dance for PD as well as (iii) before and (iv) after a matched-intensity exercise condition. All 7 subjects completed self-reported questionnaires on Positive and Negative Affect (PANAS-X) and on Body Self-Efficacy (BSE) at these four time points. Only 5 subjects completed the set of four physiological recordings (i.e., heart rate and electrodermal activity), as well as 5 subjects successfully completed four assessments of gait and dual task performance, due to subjects’ delays, device failures and/or malfunction.

### 2.3 Participants

Subjects’ baseline data were collected during an initial assessment, including PD duration, age, Hoehn & Yahr stage, and Movement Disorder Society (MDS) Unified Parkinson’s Disease Rating Scale (UPDRS) scores (Table 1). The MDS-UPDRS scores comprise four sections: 1-Non-Motor Aspects of Experiences of Daily Living (1A-Complex behaviors (6 items); 1B-Patient Questionnaire (7 items)); 2-Motor Aspects of Experiences of Daily Living (13 items); 3-Motor Examination (33 items); 4-Complications of Therapy (6 items). All MDS-UPDRS scores were collected when subjects were ON their regular medication schedule. In one case, subject 7 was tested during OFF time, a functional state associated with the medication wearing off. Unfortunately, it was not possible to repeat the UPDRS 3 assessment, but it should be taken into account that the UPDRS 3 score may not reflect the subject’s best motor performance. Table 2 shows the medication list for all subjects with the exception of subject 3, who was the most recently diagnosed, although not the youngest, and the only subject not currently treated with levodopa, dopamine agonists or other dopamine-interacting drugs. Individuals who had PD for a longer time (subject 1, 2, and 5) had higher MDS-UPDRS 3 scores, thus more pronounced motor symptoms (Table 1). However, subject 4 demonstrated the highest MDS-UPDRS 3 score despite taking the highest medication dose relative to the other study participants (Table 2) and having had a relatively shorter disease duration.

**Table 1.** Demographic and clinical characteristics of the Dance for PD research group. Age was calculated as (date of assessment-date of birth)/362.25. Hoehn & Yahr is reported to indicate the stage of disease progression (0-5). MDS-UPDRS scores include parts I: Non-Motor Aspects of Experiences of Daily Living (IA, Complex behaviors (6 items); IB, Patient Questionnaire (7 items)); II: Motor Aspects of Experiences of Daily Living (13 items); III: Motor Examination (33 items); IV: Complications of Therapy (6 items). SD: Standard Deviation.

| Subject ID | PD duration (years) | Age (years) | Hoehn & Yahr | UPDRS IA | UPDRS IB | UPDRS II | UPDRS III | UPDRS IV |
| --- | --- | --- | --- | --- | --- | --- | --- | --- |
| 1 | 9.7 | 59.5 | 2 | 2 | 10 | 7 | 24 | 4 |
| 2 | 8.9 | 75.1 | 2 | 5 | 8 | 9 | 19 | 7 |
| 3 | 1.5 | 74.5 | 1 | 2 | 10 | 5 | 9 | 0 |
| 4 | 4.2 | 67.5 | 2 | 11 | 6 | 7 | 34 | 0 |
| 5 | 9.8 | 69.1 | 2 | 1 | 14 | 6 | 25 | 4 |
| 6 | 3.0 | 80.3 | 1 | 1 | 2 | 2 | 6 | 0 |
| 7 | 3.6 | 74.0 | 1 | 13 | 7 | 4 | 20 | 1 |
| **Mean** | **5.8** | **71.4** | **1.6** | **5.0** | **8.1** | **5.7** | **19.6** | **2.3** |
| **SD** | **3.5** | **6.7** | **0.5** | **5.0** | **3.8** | **2.3** | **9.6** | **2.8** |

**Table 2.**
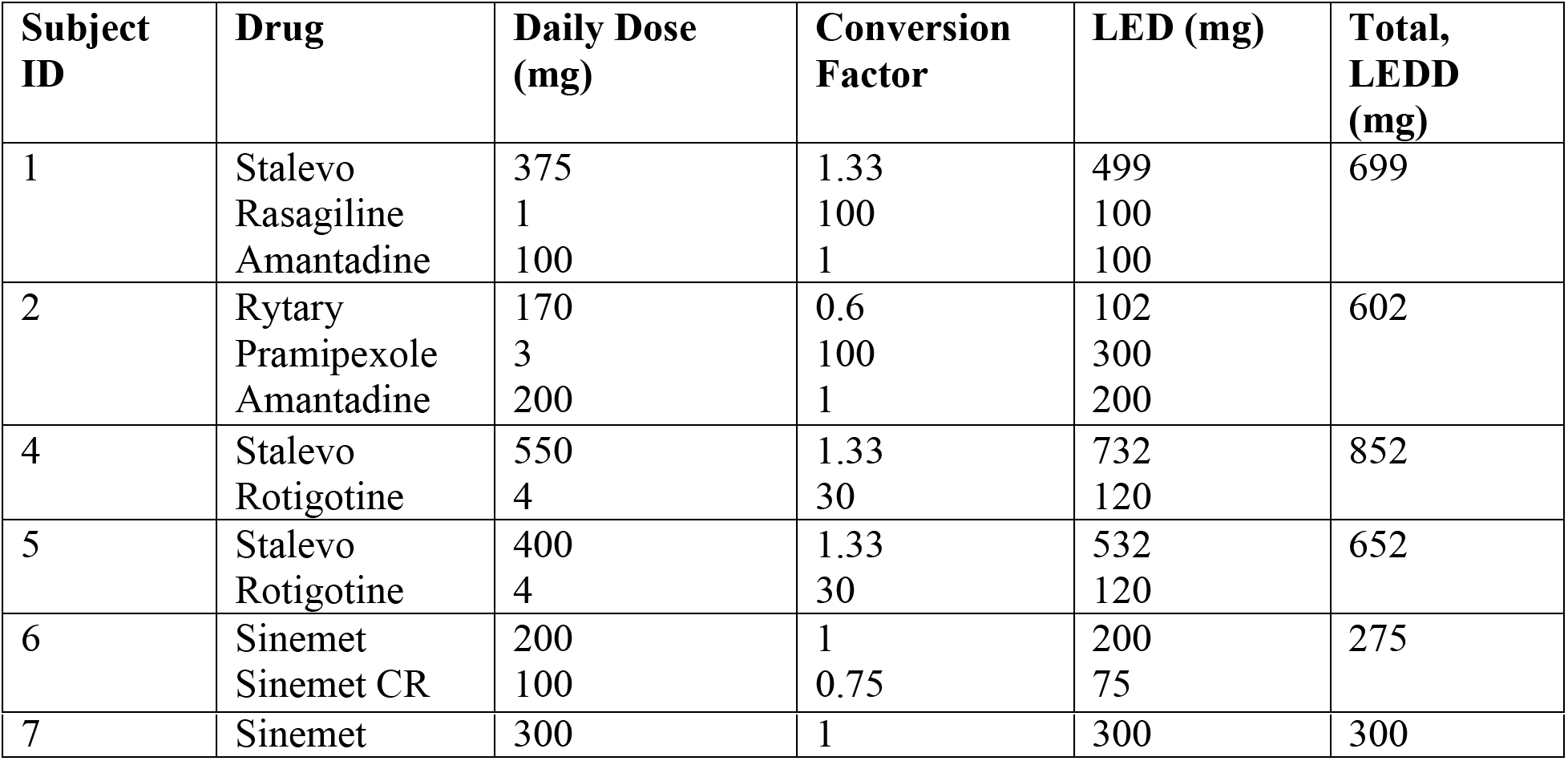
Levodopa equivalent daily dose (LEDD) calculation. For drugs containing levodopa and other active components (e.g., carbidopa), we report the levodopa amount only. We also show the conversion factors that lead to the LED calculation following Tomlinson (2010) guidelines. Subject 3 is not reported since not currently treated with levodopa, dopamine agonists or other dopamine-interacting drugs.

Since this study investigated affective, self-efficacy related, and motor changes in response to two different movement experiences (Dance for PD, matched-intensity exercise), other relevant baseline characteristics were collected, including depression, cognitive, and gait scores (Table 3).

**Table 3.** Other Clinical scores of the Dance for PD research group. Beck Depression Inventory-II (BDI-II) (21 items), Montreal Cognitive Assessment (MoCA), Gait and Falls Questionnaire (GFQ) (16 items), Freezing of Gait score (FoG) (6 out of 16 GFQ items). SD: Standard Deviation.

| Subject ID | BDI-II | MoCA | GFQ | FoG |
| --- | --- | --- | --- | --- |
| 1 | 6 | 26 | 15 | 8 |
| 2 | 8 | 25 | 12 | 10 |
| 3 | 7 | 26 | 1 | 0 |
| 4 | 6 | 30 | 6 | 2 |
| 5 | 12 | 27 | 4 | 2 |
| 6 | 2 | 27 | 1 | 0 |
| 7 | 14 | 24 | 3 | 1 |
| **Mean** | **7.9** | **26.4** | **6.0** | **3.3** |
| **SD** | **4.0** | **1.9** | **5.5** | **4.0** |

Depressive symptoms were assessed using the Beck Depression Inventory-II (BDI-II) (Beck et al., 1996). Individual scores indicate minimal depression (BDI-II score 0-13) with only one subject in the range of mild depression (BDI-II score 14-19). The Montreal Cognitive Assessment (MoCA) was used to screen the participants for mild cognitive impairment (MCI), as this tool is more sensitive than the Mini-Mental State Examination in this population (Hoops et al., 2009). However, it has been previously suggested that lower MoCA scores in PD patients may reflect worse executive function performance rather than MCI (Chou et al., 2014), so in this study the MCI screening cutoff point was set at 24/25 rather than 26/27 (Hoops et al., 2009). The average MoCA score in this group was 26.4. (SD = 1.9), with one subject scoring 24 and every other subject above that value. Finally, the self-reported Gait and Falls Questionnaire, and the related Freezing of Gait score (Giladi et al., 2000), were collected to provide context to the subjects’ gait performance later assessed through the 6-minute walking test (6MWT) and the Timed-Up-and-Go test (TUG).

### 2.4 Dance for PD and Matched-Intensity Exercise

Our hypothesis is that a Dance for PD class would have different effects than a matched-intensity exercise session lacking artistic elements like the presence of music, the use of narrative and metaphorical language, and a social reality of grace and beauty, established and reinforced by dance teachers, live musicians, and a group of peers with PD addressed as “dancers.” These elements have been identified in light of the embodied aesthetics model (Koch, 2017) which defines beauty, imagery, symbolization, meaning making, self-efficacy, and community as active therapeutic factors of the art therapies.

To test this hypothesis, the researcher, who is a certified movement analyst (CMA), designed an ad-hoc control intervention to compare, within the same subjects, a Dance for PD class to a movement intervention both presented at the same location and time of the day, while also matching aerobic intensity, class structure (e.g., from upper body to lower body, from sitting to standing), class progression from single movements to sequences of increasing complexity, and attention to body organization, spatial directions, and rhythm. Participants were scheduled on two different days within the same week. On one day, the participants were tested before and after a regular Dance for PD class. On a different day, within the same week, testing was conducted before and after the matched-intensity exercise session. Both classes took place between 2:15 pm and 3:30 pm at The Mark Morris Dance Center in Brooklyn, NY. All participants were tested before and after both Dance for PD and matched-intensity exercise. However, the order of these two conditions could not be fully balanced due to the participants schedule and availability. In particular, subjects 2 and 3 scheduled the matched-intensity exercise session two days before the Dance for PD class, while subjects 1, 4, 5, 6, and 7 returned two days after the Dance for PD class for the matched-intensity exercise condition and testing. The same experimenter met people at the class, consented, tested before and after both Dance for PD and matched-intensity exercise.

Dance for PD classes were taught by a certified instructor following a specific structure, progressing from a seated warm-up to seated dynamic sequences. The classes develop through (chair or barre) supported standing combinations and lead up to traveling in space through the dance studio. Core values of these classes include artistic excellence (the classes are taught by professional dancers and are accompanied by live musicians), creative expression (movement combinations include narrative elements, metaphorical language, and musicality), and community engagement (participants are addressed as dancers and are invited to move and connect through shared gesture and movement).

The matched-intensity exercise was designed to parallel the structure of a Dance for PD class, mobilizing upper and lower joints while sitting on a chair, then transitioning to standing and finally walking in space. Importantly, the intensity of both dance and exercise was assessed by Heart Rate (HR) measurements, and Percentage Heart Rate Reserve (PHRR) calculation, to compare the physical activity intensity levels of these sessions (Ignaszewski et a., 2017). Table 4 reports the four-parts structure of the class (sitting upper body, sitting lower body, standing, and walking), each part consisting of different movements focused on joints and limbs mobilization. Single movements (e.g., spinal flexion) were initially presented and experienced through repetition, while later composed in sequences of increased complexity (i.e., series of different movements in succession). The subjects were welcomed into a dance studio, set up at a table with a chair, and introduced to the computer set up (Table 4). All participants completed the movement sessions following the prompts from a computer screen in a dance studio at Mark Morris Dance Center, the same setting as the Dance for PD classes. Importantly, it was necessary to remove professional dancers and musicians from the matched-intensity exercise condition, since the scope of this study was to investigate whether the artistic context of the movement experience could mediate some of the effects of the dance intervention. In order to manipulate this interpersonal (i.e., social) context, the matched-intensity exercise class was presented through both written language and images on a computer screen, a modality that matches exercise and fitness programs prescribed to people with PD at home. The instructions provided told participants which movement to perform (e.g., “turn”), which body part to engage (e.g., “the head”), in which spatial direction (e.g., “to the right”), the number of repetitions involved (e.g., “you will repeat 10 times”), and at which tempo to move through the beat provided by a metronome associated with the images presented. The images included both stylized and real human bodies, drawn from the online UK National Health Service (2018) physical activity guidelines for adults as well as from Bezner and Rose (1989) adult exercise instruction sheets. These images are sources that people could access to work out at home, as well as exercise routines frequently used by physical and occupational therapists in older adult rehabilitation settings.

**Table 4.** Structure and presentation of the matched-intensity exercise condition. The class included four parts (sitting - upper body, sitting - lower body, standing, walking), each part consisting of different movements focused on joints and limbs mobilization. Single movements were initially presented and later composed in sequences of increased complexity (i.e., series of single movements in succession).

| Class Structure | Movement |
| --- | --- |
| Sitting - Upper Body | Flexion and extension of the spine<br>Rotation and lateral flexion of the neck<br>Elevation, adduction and abduction of the scapula<br>Circumduction of the arms<br>Flexion and extension of the elbows<br>Circumduction of the wrists |
| Sitting - Lower Body | Abduction and adduction of the hips<br>Medial and lateral rotation of the hips<br>Hip flexion<br>Flexion and extension of knees<br>Circumduction of the ankles<br>Dorsiflexion and plantar flexion of the feet |
| Standing | Standing up from sitting<br>Flexion and extension of knees<br>Extension of the toes<br>Abduction of one leg<br>Extension of hips<br>Lateral flexion of spine |
| Walking | Single side steps<br>Locomotion |

### 2.5 Outcome Measures

#### 2.5.1 Heart Rate

Heart rate was measured during both Dance for PD and matched-intensity exercise using the E4 wristband, a wearable device that can be placed on either wrist to record physiological and motion features for a prolonged time (Empatica, 2018). The wristband includes four sensors for data acquisition, providing photoplethysmogram (PPG), electrodermal activity (EDA), 3-axial movement, and skin temperature (Garbarino et al., 2014). The PPG sensor has a non-customizable sampling frequency of 64 Hz and employs an artifact removal technique that operates through a combination of green and red light. These multiple wavelengths ensure accuracy when the wrist activity observed though accelerometers are not reliable or significant (Empatica, 2018). The PPG sensor provides the blood volume pulse (BVP) from which heart rate (HR) was derived. To analyze the E4 data, the Toolbox for Emotion Analysis using Physiological signals (TEAP) was employed in Matlab (Version 9.5.0.944444, R2018b, The Mathworks, Inc., Natick, MA) enabling pre-processing and extraction of features from the PPG signal (Soleymani et al., 2017). After PPG data were imported, the BVP signal was acquired, and segments of 5 minutes (rest) and 60 minutes (activity) were selected from the recording. The BVP signal was smoothed using a moving median with a 2 second window (i.e., 128-time points), to remove artifacts such as rapid shocks or other anomalies (Peper et al., 2010).

Consequently, the HR signal was extracted using the TEAP function “*BVP_feat_BPM*”, in units of beats per min (BPM). From the HR signal, HR means were computed at rest and during activity for both the Dance for PD and matched-intensity exercise conditions. In order to compare the level of physical activity during the two conditions, we computed the Percentage Heart Rate Reserve (PHRR), as the ratio of the difference between HR at rest (HR_rest_) and HR during physical activity (HR_act_) to the difference between HR at rest (HR_rest_) and maximum HR (HR_max_). HR_max_ was predicted using Tanaka’s formula 208 - (0.7 × age) (de Oliveira Segundo et al., 2016).

#### 2.5.2 Electrodermal Activity

The E4 wristband is equipped with an Electrodermal Activity (EDA) sensor designed to measure the electrical conductance of the skin in the ventral area of the wrist, with two stainless steel dry electrodes that apply a small alternating current and measure the resulting current flow between them. The EDA is measured in micro Siemens (μS) and sampled at a frequency of 4 Hz (Empatica, 2018). In general, the activity of sweat glands, innervated by the sympathetic nervous system, determines the skin conductance values (Critchley, 2002). EDA analysis includes two components, the overall skin conductance level (SCL), which is the tonic level of electrical conductivity of the skin, and the skin conductance response (SCR), which comprises the phasic changes in electrical conductivity that may occur whether or not an identifiable stimulus is provided (Braithwaite et al., 2015). The data was imported in Matlab (Version 9.5.0.944444, R2018b, The Mathworks, Inc., Natick, MA) and the raw EDA signal was cut to 60 minutes of activity, during either Dance for PD or matched-intensity exercise. After applying a moving median smoothing to the raw EDA signal (2 seconds window, i.e. 8-time points), the tonic features, including the overall skin conductance level (SCL) and the amplitude of spontaneous fluctuations (SF) were computed. The spontaneous fluctuations were derived by defining peaks in the signal as amplitude increases greater than 0.05 μS over an interval of 5 seconds or less (Braithwaite et al., 2015).

#### 2.5.3 Affect, Self-Efficacy, Beauty

The Positive and Negative Affect Schedule (PANAS-X) is a self-reported questionnaire that can be completed in 8-10 minutes by most subjects (Watson and Clark,1994). It comprises 60-items, terms describing different moods, to provide measurements of two general dimensions of affect (General Negative Affect, GNA; General Positive Affect, GPA), as well as four complex affective states (shyness, fatigue, serenity, and surprise). In this study, the subjects were instructed to indicate the extent to which they felt a specific mood in the present moment, using a 5-point Likert scale to report their answer (1 = very slightly or not at all, 2 = a little, 3 = moderately, 4 = quite a bit, 5 = extremely). The PANAS-X was administered both before and after the Dance for PD class as well as before and after the matched-intensity exercise condition. In tandem with this questionnaire, the subjects were asked to fill in another self-reported questionnaire on Body Self Efficacy (BSE) (Fuchs and Koch, 2014). This 10-item scale measures the subjective perception of one own bodily ability (e.g., “I can move well” or “I can express myself in movement” vs “I have many bodily constraints” or “My body is lifeless and inert/numb”). The subjects were asked to answer how each statement applied to them in the present moment, rating their responses from zero to five (0 = does not apply at all, 5 = applies exactly). The items include two accounts that constitute a sub-dimension on aesthetic experience, which state “My movements are beautiful” and “I can move elegantly/with grace” (Koch et al., 2016). Four reports were collected for each subject, both before and after Dance as well as before and after matched-intensity exercise.

#### 2.5.4 Gait performance during the 6-minute walking test

G-WALK is a portable system for functional assessment of movement, produced by BTS Bioengineering (2016). This electronic device is also called an inertial measurement unit (IMU), which provides quantitative data reports about acceleration, angular rate, and body orientation through a combination of a) 3-axial accelerometers, a combination of three linear accelerometers measuring acceleration along the x, y, and z-axis respectively; b) gyroscopes, sensing angular velocity along each rotational axis; and c) magnetometers; providing the direction or heading of the movement with respect to the Earth’s magnetic field.

The G-WALK inertial system can perform and analyze frequently used clinical motor tests, two of which are particularly relevant to PD patients’ assessment: the 6-minute walking test (6MWT) to measure a patient’s functional capacity (Bautmans et al., 2004), and the timed-up-and-go (TUG), for balance, mobility and risk of fall assessment (Podsiadlo and Richardson, 1991; Shumway-Cook et al., 2000). The device is intended to assess patients’ progress between admission and discharge or to evaluate the degree of functional improvement that can be achieved with a therapy session or a specific orthotic device.

To perform these tests, the sensor was connected via Bluetooth to the G-Studio software on a PC, inserted into a belt, and placed in the back of the subject. Importantly, each test requires a specific sensor position to ensure reliable data collection. In particular, the 6MWT is performed by placing the sensor at S1-S2 level, the transition between lumbar and sacral vertebrae, while the TUG requires sensor at L2 level, right below the transition between thoracic and lumbar vertebrae (i.e., under the superior lumbar L1 vertebrae).

All subjects were able to walk without physical assistance or assistive devices. To collect the 6MWT data, the subjects begun by standing still, allowing for the device stabilization, then started to walk at a comfortable pace for six consecutive minutes, along a 10-meters linear path, turning 180° at the two peripheral points. When the test terminated, at the end of the six minutes, the performance report was saved. The subjects repeated this test before and after Dance for PD, as well as before and after the matched-intensity exercise condition, generating four reports for each subject. The 6MWT reports provide values of velocity (m/s) as the average walking speed, cadence (steps/min) as the average number of steps per minute, stride cycle length (m) as the average distance covered from one initial contact to the next on the same side, as well as the symmetry index (%) which provides a comparison of the anteroposterior acceleration between the right and left gait cycles.

#### 2.5.5 Dual Task Performance with the Timed-Up-and-Go test

The TUG test is used as a tool for the assessment of lower limb mobility and function, as well as fall risk, with subjects walking at their normal pace (Podsiadlo and Richardson, 1991). A modified version of the TUG test including a cognitive task (TUG-cog) was introduced to better predict the fall risk in older adults (Shumway-Cook et al., 2000). In the TUG-cog test, subjects are instructed to count backward by threes from a random number between 60 and 100.

To perform the TUG test, the subject started in a seated position, with both arms relaxed on the thighs. The device stabilization phase took place with the subject seated in a static position and relaxed. From this resting position, the subjects were instructed to stand up, walk 3 meters, turn 180° walk back to the seat, and sit down again. A piece of tape was placed on the floor at 3 meters to be easily seen by the subject as reference. Participants were tested before and after Dance for PD as well as before and after the matched-intensity exercise condition. Since the subjects were wearing their regular footwear, they were asked to be consistent in the two testing days. At each of these four time-points, they were given a practice trial to rehearse the movement, followed by two formal trials for the TUG test and two formal trials for the TUG-cog test. The average of the two trials was analyzed for both TUG and TUG-cog.

We calculated an average score for single (TUG test) and dual task (TUG-cog), and then computed the dual task cost (in seconds), which represents the extra time needed for the dual task performance. Finally, we calculated the difference in dual task cost both before and after Dance for PD, as well as before and after matched-intensity exercise.

### 2.6 Statistical Analysis

The comparison between Dance for PD and matched-intensity exercise was based on matched pairs (every subject was tested for both conditions) leading to a series of differences (i.e., before-after Dance for PD, before-after matched-intensity exercise). The Wilcoxon Rank-Sum test for paired samples was employed to run the comparisons across conditions of the before-after differences in Dance for PD and matched-intensity exercise, respectively. A sample of 5 subjects was analyzed for both heart rate and electrodermal activity, while 7 subjects’ self-reported questionnaires on affect and self-efficacy were examined. Also, 5 subjects were analyzed for the 6-minute walking test (6MWT), as well as 5 subjects for the Timed-Up-and-Go test (TUG) with cognitive task (TUG-cog). Importantly, The Wilcoxon Rank-Sum test is a non-parametric test that allows the comparison between two related-groups (Dance for PD, matched-intensity exercise) with no assumption of distribution or normality. Thus, the Wilcoxon Rank-Sum test allows comparison between the before-after differences in Dance for PD and in matched-intensity exercise, offering a non-parametric alternative to repeated measures ANOVA when the statistical assumptions cannot be met due to sample size (Oberfeld and Franke, 2013).

A correlation between electrodermal activity and general positive affect was calculated using Spearman (ranked data) rather than Pearson (continuous data). Both statistics, Wilcoxon Rank-Sum test and Spearman correlation, were run on SPSS (IBM Corp. Released 2011. IBM SPSS Statistics for Windows, Version 20.0. Armonk, NY: IBM Corp.).

## 3 RESULTS

### 3.1 Heart Rate (HR) and Electrodermal Activity (EDA)

The HR signal during 60 minutes of activity during Dance for PD and matched-intensity exercise, for the five subjects recorded (subject 3,4,5,6,7) is plotted in Figure 1. HR values are summarized in Table 5, reporting the average Heart Rate (HRact) during activity, as well as the calculated percentage heart rate reserve (PHRR), during both conditions. The PHRR values are below 30%, considered within the range of “very light” physical activity (Ignaszewski et a., 2017). Importantly, the PHHR values are not significantly different between Dance for PD and matched-intensity exercise (Z = −0.135, p = 0.893), as shown in Figure 2A.

**Figure 1:**
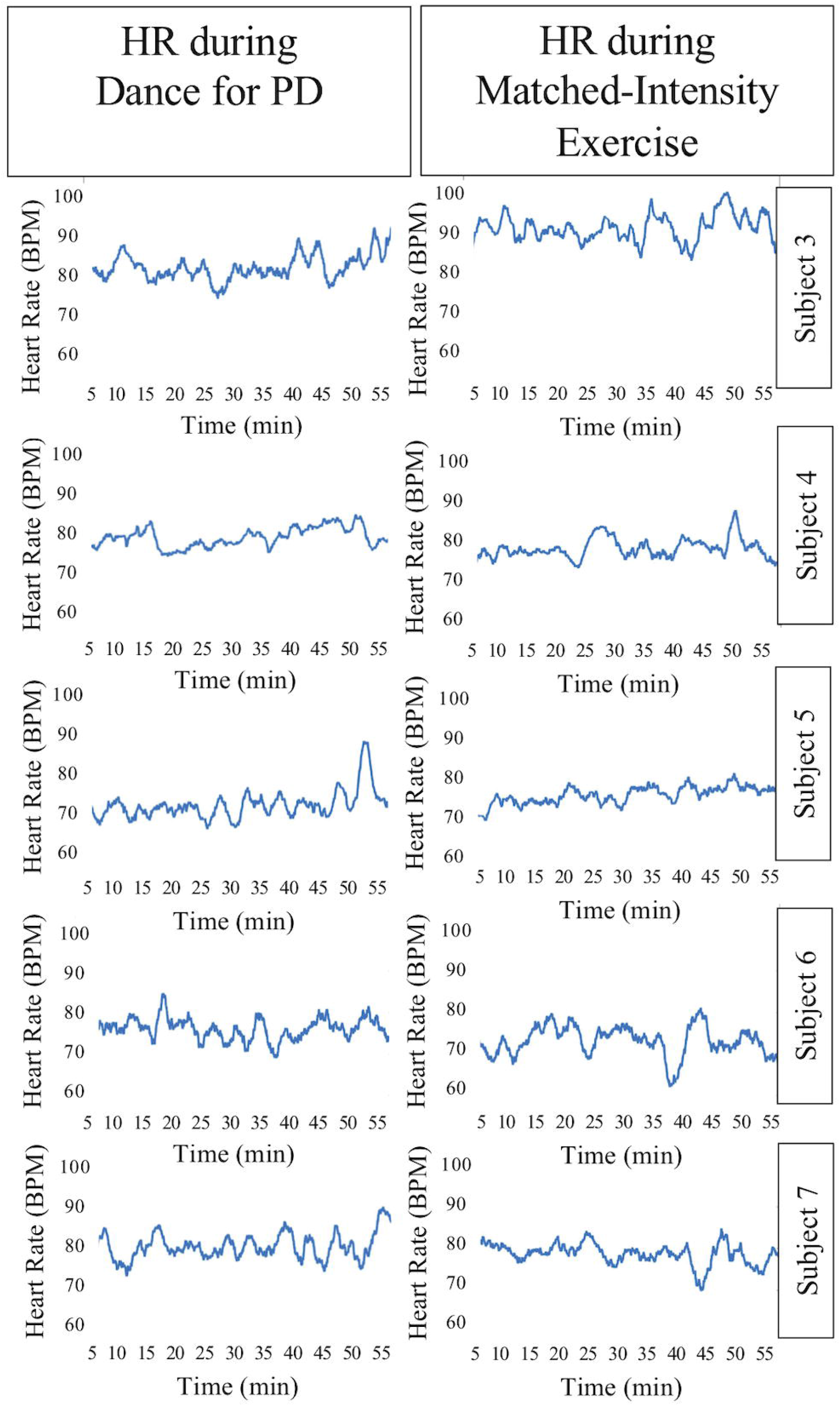
Heart Rate (HR) signal during 60 minutes of Dance for PD and matched-intensity exercise. BPM: beats per minute.

**Figure 2:**
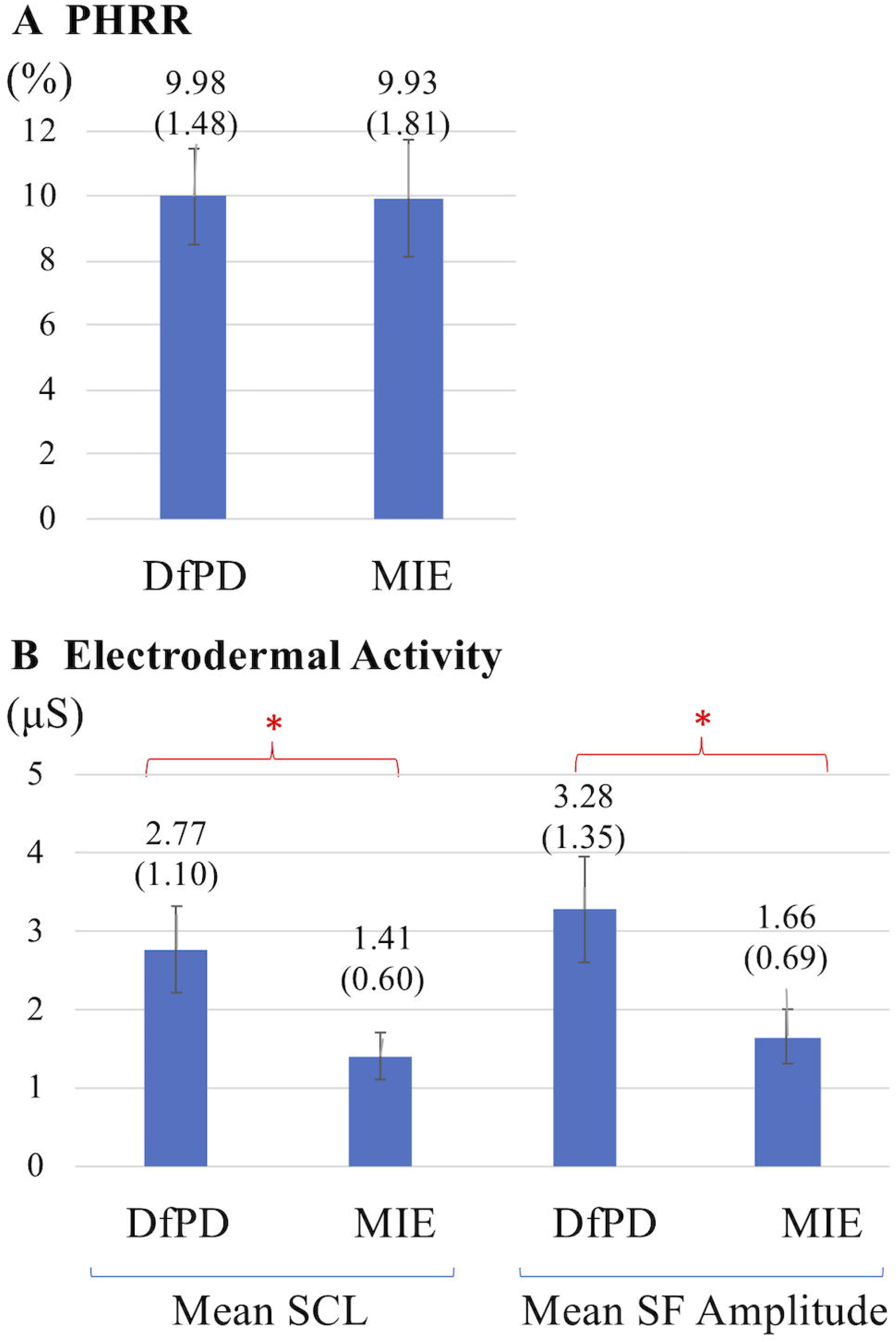
**(A)** Percentage Heart Rate Reserve (PHRR) recorded during Dance for PD (DfPD) and matched-intensity exercise (MIE). Average PHRR values are reported for both conditions with standard errors in parenthesis. **(B)** Electrodermal activity (Skin Conductance Level, SCL; Spontaneous Fluctuations, SF) during Dance for PD (DfPD) and matched-intensity exercise (MIE). Average SCL and SF values are reported for both conditions with standard error in parenthesis and error bars.

**Table 5.**
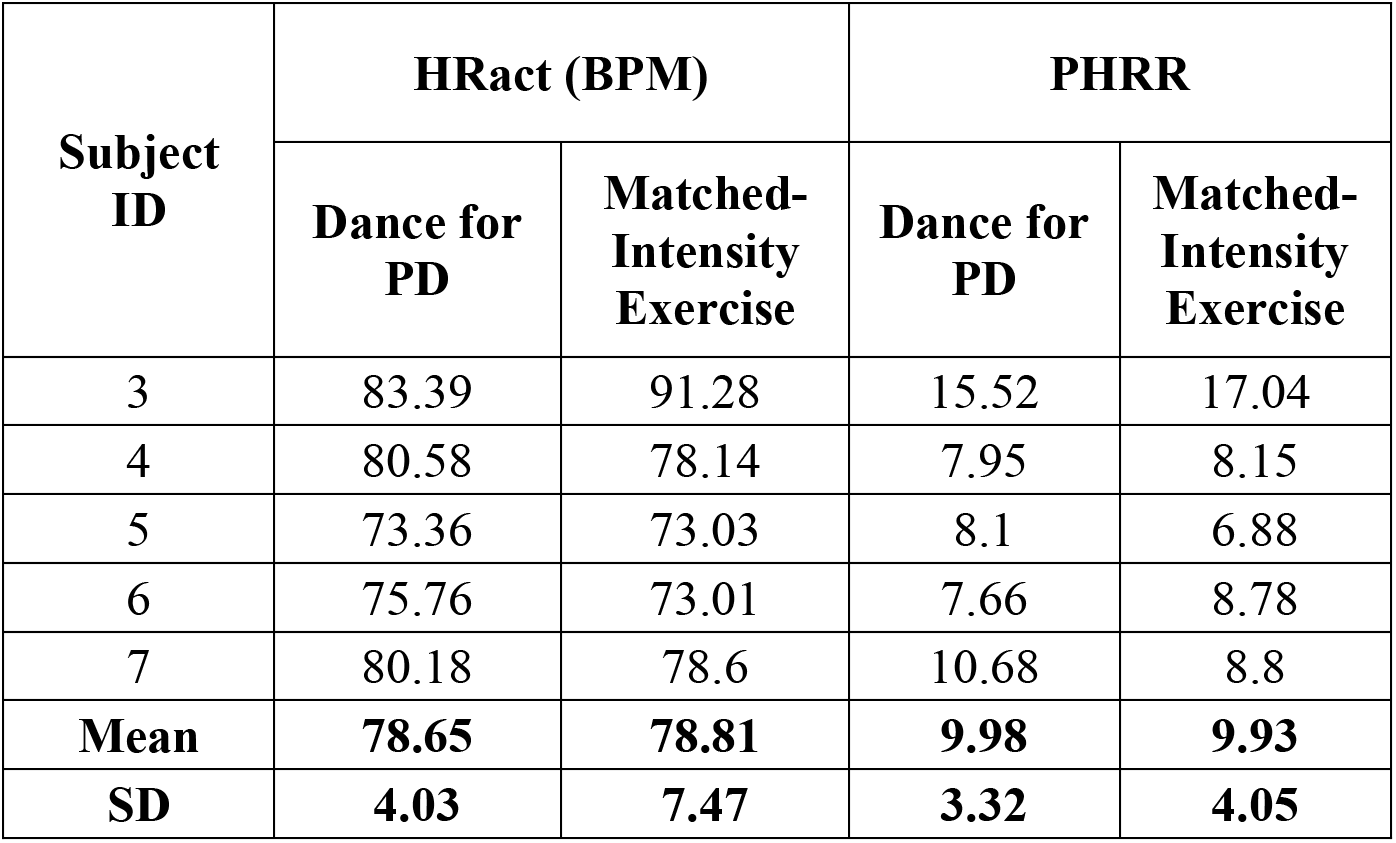
Average Heart Rate (HRact) and Percentage Heart Rate Reserve (PHRR) during Dance for PD and matched-intensity exercise for the five subjects who completed physiological recordings. BPM: beats per minute; SD: Standard Deviation.

Table 6 reports tonic electrodermal features (i.e., skin conductance level, spontaneous fluctuations) computed from the EDA signal during both conditions (i.e., Dance for PD and matched-intensity exercise) for five subjects. SCL mean amplitudes correspond to the sample mean of the skin conductance signal (in μS) over 60 minutes. The mean amplitude of spontaneous fluctuations (SF) are calculated from the peaks in the signal that occur over the same amount of time. The mean SCL (Z = −2.023, p = 0.043) and SF amplitudes (Z = −2.032, p = 0.042) are significantly higher during Dance for PD than during the matched-intensity exercise condition, as reported in Figure 2B.

**Table 6.** Mean Skin Conductance Level (SCL) and Spontaneous Fluctuations (SF) mean amplitude for both Dance and matched-intensity exercise for the five subjects who completed physiological recordings. SD: Standard Deviation.

| Subject ID | Mean SCL (μS) | | Mean SF Amplitude (μS) | |
| --- | --- | --- | --- | --- |
|  | Dance for PD | Matched-Intensity Exercise | Dance for PD | Matched-Intensity Exercise |
| 3 | 6.61 | 3.68 | 8.2 | 4.27 |
| 4 | 3.62 | 1.02 | 3.96 | 1.14 |
| 5 | 2.22 | 1.46 | 2.34 | 1.8 |
| 6 | 0.51 | 0.43 | 0.64 | 0.54 |
| 7 | 0.89 | 0.48 | 1.27 | 0.56 |
| **Mean** | **2.77** | **1.41** | **3.28** | **1.66** |
| **SD** | **2.47** | **1.34** | **3.02** | **1.55** |

### 3.2 Affect, Self-Efficacy, Experience of Beauty

Figure 3 shows the differences in self-reported PANAS-X scores for the two general dimensions of affect (General Negative Affect, GNA; General Positive Affect, GPA) and other four complex affective states (shyness, fatigue, serenity, and surprise). The item composition for these affective constructs is reported in Table 7. The differences are calculated before and after Dance for PD and matched-intensity exercise, respectively. The average trend in GPA after Dance for PD is higher than after matched-intensity exercise (Z = −1.873, p = 0.061 N.S.). The next closest trends were including surprise (Z = −1.476, p = 0.140 N.S.), GNA (Z = −1.265, p = 0.206 N.S.), and shyness (Z = −0.962, p = 0.336 N.S.) but all failed to reach significance. For the five subjects who completed both all PANAS-X self-reporting and electrodermal activity recordings, Figure 4 reports the correlation between the levels of General Positive Affect (GPA) reported after Dance for PD and the average Skin Conductance Levels (SCL) recorded during the dance session (Spearman’s rho = 0.900, p = 0.037).

**Figure 3:**
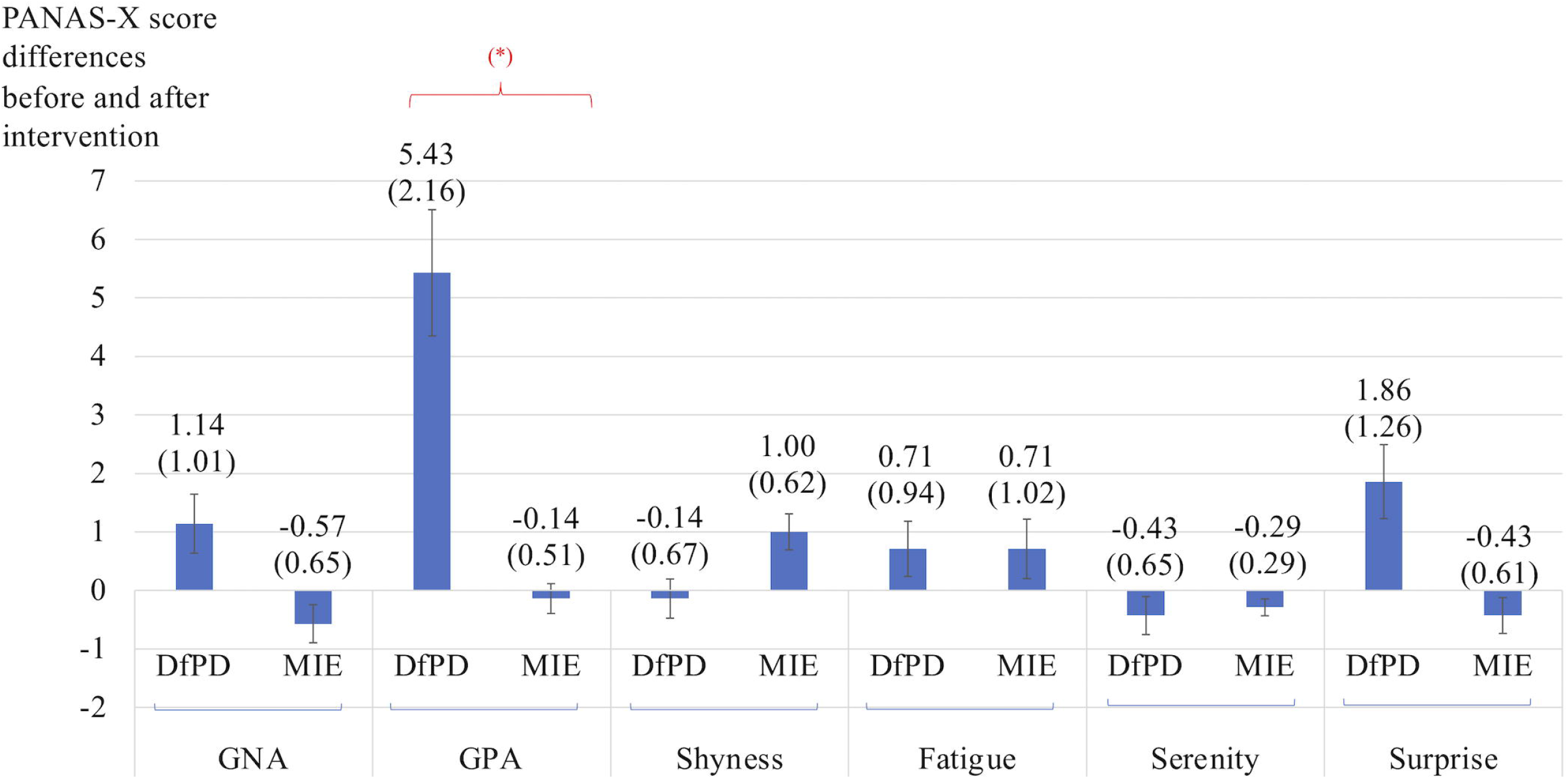
Differences in self-reported scores for the two general dimensions of affect (General Negative Affect, GNA; General Positive Affect, GPA) and other four complex affective states (shyness, fatigue, serenity, and surprise). The differences are calculated before and after Dance for PD (DfPD) and matched-intensity exercise (MIE), respectively. (*) signifies a trend at p=0.061. The PANAS-X score differences are reported as averages between seven subjects, with standard error in parenthesis and error bars.

**Figure 4:**
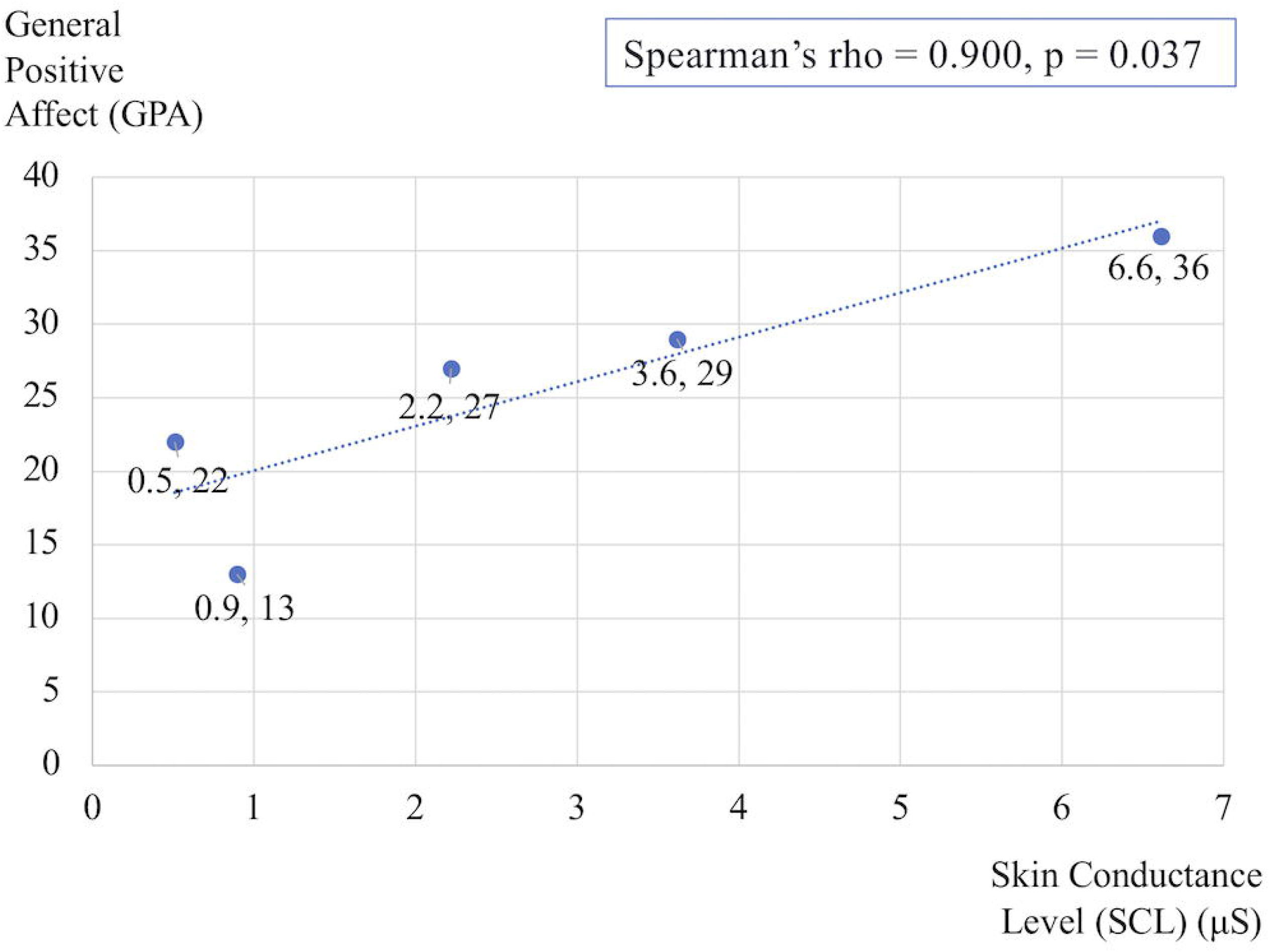
Correlation between the General Positive Affect (GPA) self-reported scores after Dance for PD and the average Skin Conductance Levels (SCL) recorded during the dance session. The data reflects changes in the five subjects who completed both PANAS-X and electrodermal recordings.

**Table 7.**
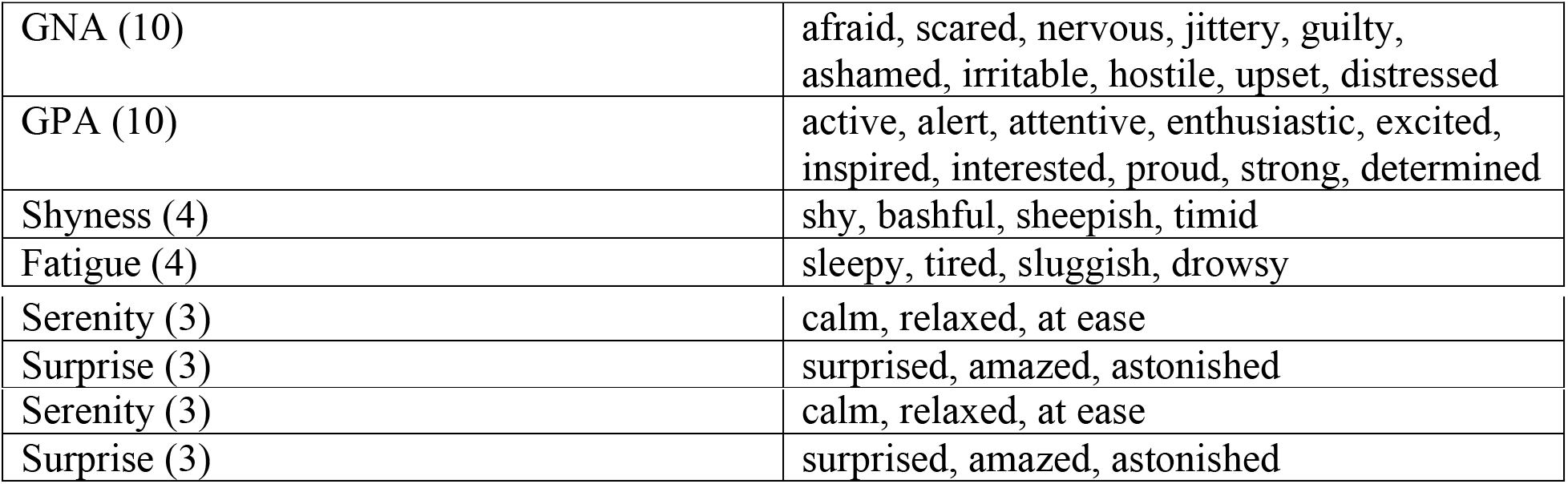
Item Composition in the PANAS-X, Manual for the Positive and Negative Affect Schedule (Watson and Clark, 1994). General Negative Affect (GNA), General Positive Affect (GPA), Shyness, Fatigue, Surprise, and Serenity. In parentheses, the number of items defining each scale.

In parallel to self-reported affect, Figure 5 shows significant differences in Body Self-Efficacy (BSE) along with changes in the beauty subscale, which focuses on 2 out of 10 BSE items. The comparison between the two interventions reveals a significant increase in both BSE scores (Z = −2.371, p = 0.018) and beauty subscale scores, (Z = −2.121, p = 0.034) after Dance for PD compared to matched-intensity exercise.

**Figure 5:**
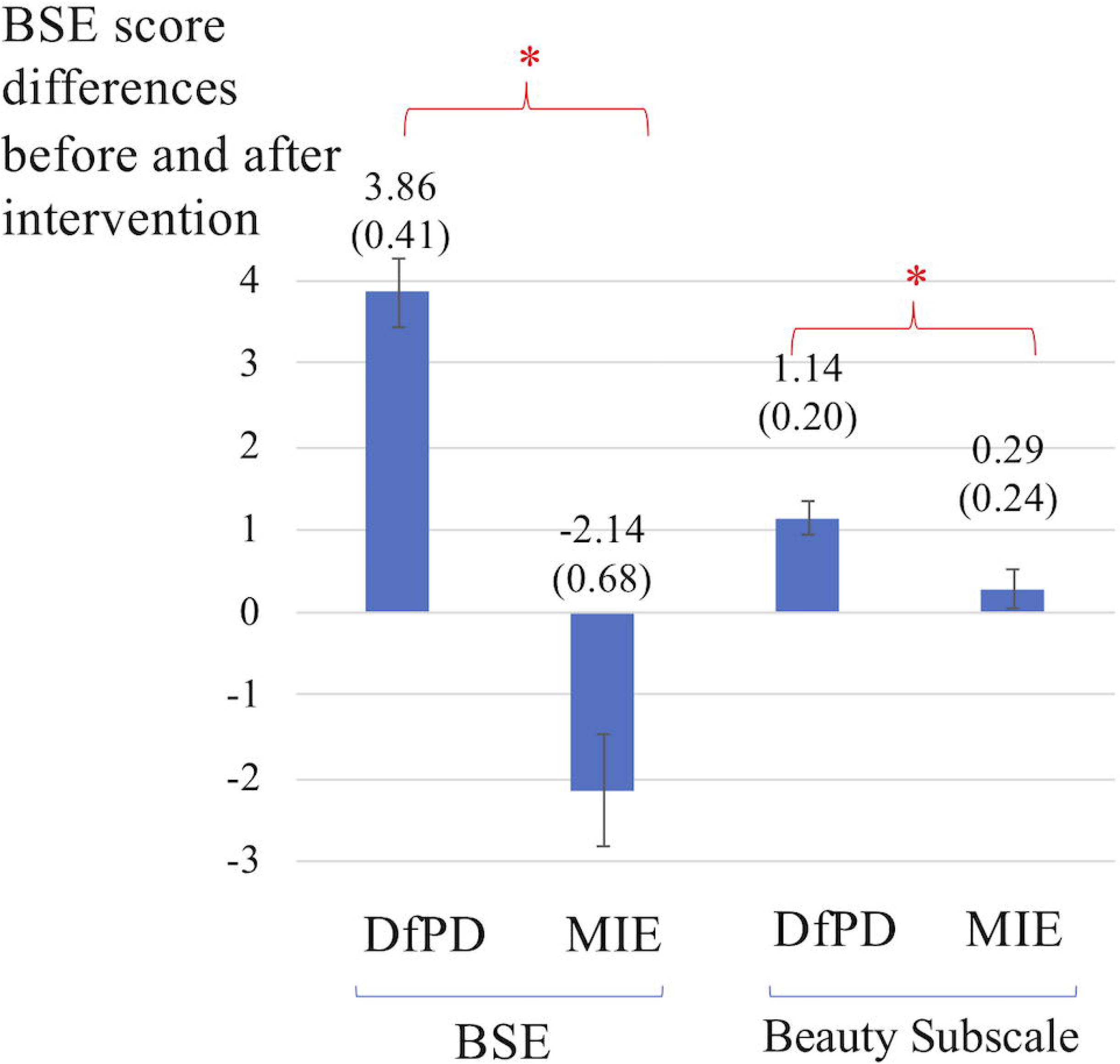
Differences in self-reported scores for Body Self-Efficacy (BSE) and the beauty subscale, which captures 2 out of 10 BSE items. The differences are calculated before and after Dance for PD (DfPD) and matched-intensity exercise (MIE), respectively. The BSE score differences are reported as averages between subjects, with standard error in parenthesis and error bars.

### 3.3 Gait performance during the 6-minute walking test (6MWT)

Changes in mean velocity (Z = −1.490, p= 0.136 N.S.), mean cadence (Z = −1.095, p= 0.273 N.S.), and mean stride cycle length (Z = −1.483, p= 0.138 N.S.) before and after Dance for PD are not different than before and after matched-intensity exercise, as reported in Table 8.

**Table 8.** Changes in Mean Velocity, Mean Cadence, and Mean Stride Cycle Length during the 6-minutes walking test (6MWT), calculated as differences before and after Dance for PD and Matched-Intensity Exercise, respectively. SD: Standard Deviation.

|  | Mean Velocity<br>(m/s) |  | Mean Cadence<br>(steps/min) |  | Stride Cycle<br>Length (m) |  |
| --- | --- | --- | --- | --- | --- | --- |
| Subject ID | Dance for PD | Matched-Intensity Exercise | Dance for PD | Matched-Intensity Exercise | Dance for PD | Matched-Intensity Exercise |
| 1 | 0.29 | 0.02 | 2.20 | -3.70 | 0.31 | 0.07 |
| 2 | -0.10 | -0.15 | -4.20 | -4.20 | -0.07 | -0.12 |
| 4 | -0.02 | 0.02 | -3.20 | -0.90 | 0.00 | 0.03 |
| 5 | 0.02 | 0.01 | 2.00 | 1.50 | 0.00 | -0.01 |
| 7 | 0.13 | -0.14 | 9.10 | 0.30 | 0.04 | -0.12 |
| **Mean** | **0.06** | **-0.05** | **1.18** | **-1.40** | **0.06** | **-0.03** |
| **SD** | **0.15** | **0.09** | **5.30** | **2.48** | **0.15** | **0.09** |

Most importantly, Figure 6A illustrates the increases in symmetry index (%) values after Dance for PD, in front of the decreases in symmetry index values after matched-intensity exercise, for each subject. Figure 6B shows the statistically significant difference (Z = −2.032, p= 0.042) between the average increase in symmetry index (%) across subjects following Dance for PD and the average decrease across subjects following matched-intensity exercise.

**Figure 6:**
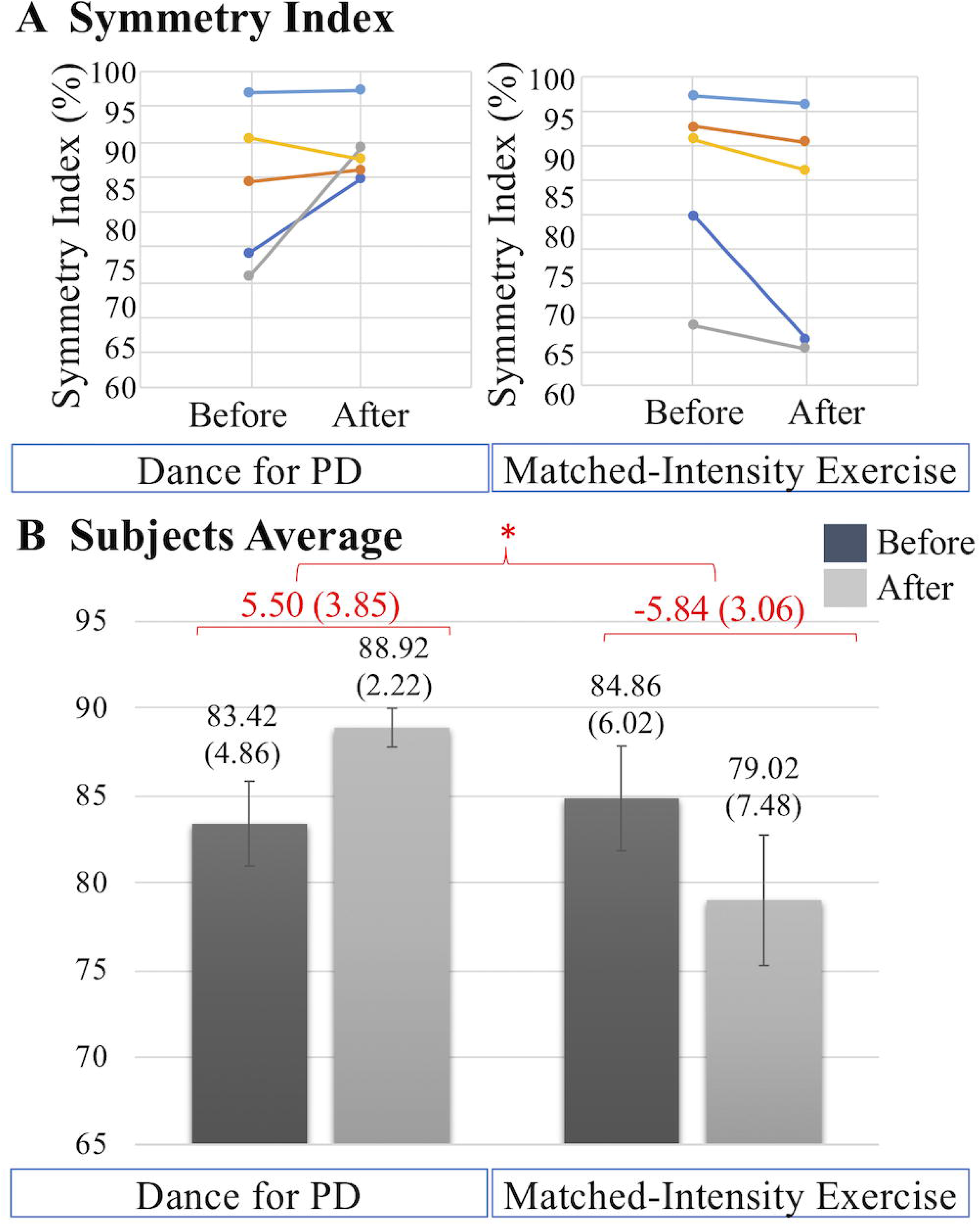
**(A)** Symmetry index (%) values before and after Dance for PD and matched-intensity exercise in five subjects (1 dark blue, 2 orange, 4 grey, 5 yellow, 7 light blue). Increases in symmetry index indicate gait improvement. **(B)** Averages across subjects of symmetry index values before and after Dance for PD, as well as before and after matched-intensity exercise. Mean values are reported, error bars indicate standard error (also in parenthesis). In red, the means across subjects of the differences between before and after, for both Dance for PD and matched-intensity exercise, with standard error in parenthesis.

### 3.4 Dual Task Performance with the Timed-Up-and-Go test (TUG)

Table S1 (Supplementary Material) reports negative values representing decreases in dual task cost, which are decreases in the slowing down of the motor performance due to the concurrent cognitive task, defining a more efficient TUG performance. These values are reported separately for each of the following sub-phases of the Timed-Up-and-Go (TUG) test: sit-to-stand, delay before forward gait, forward gait, mid turning, return gait, and end turning with stand to sit (computed together). The decreases in dual task cost in each single subphase are not significantly larger after Dance for PD compared to the matched-intensity exercise, including the sit-to-stand phase (Z = −1.753, p = 0.080), delay before the forward gait phase (Z = −1.753, p = 0.080), forward gait (Z = −1.214, p = 0.225), mid turning (Z =-1.483, p = 0.138), return gait (Z = −1.405, p = 0.686), and end turning & stand-to-sit (Z = −1.214, p = 0.225).

However, the overall decrease in dual task cost is significantly different following the two interventions (Z = −2.023, p= 0.043). Figure 7A illustrates the decreases in dual task cost without distinction between subphases. The decreases in dual task cost after Dance for PD are more pronounced than after matched-intensity exercise. Figure 7B reports the dual task cost averages across subjects before and after the two conditions, showing a larger decrease after Dance for PD than matched-intensity exercise. The before and after differences, across subjects, representing the decreases in dual task cost, are significantly larger after Dance for PD than after matched-intensity exercise.

**Figure 7:**
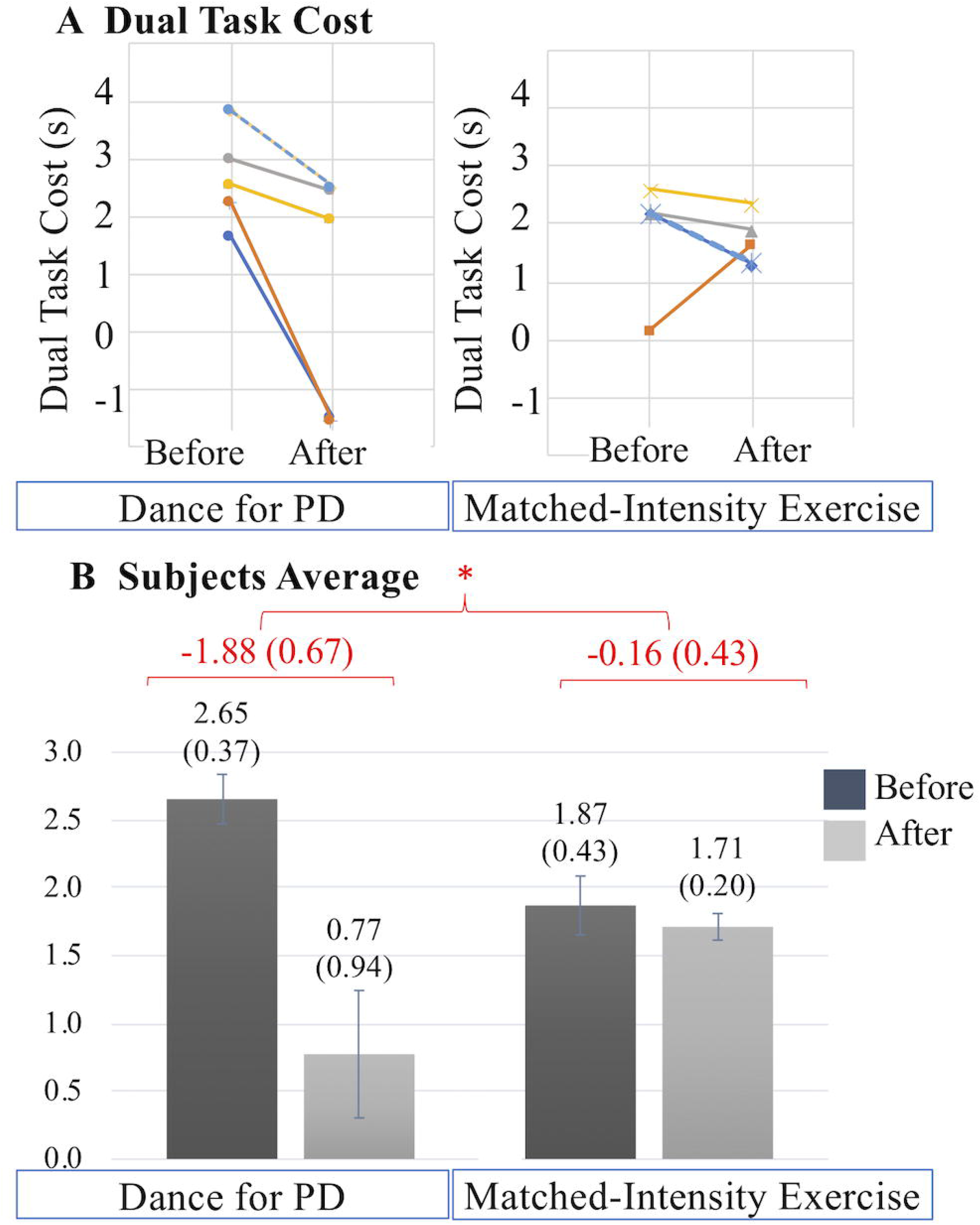
**(A)** Dual task cost (s) values before and after Dance for PD and matched-intensity exercise in five subjects (2 dark blue, 3 orange, 4 grey, 5 yellow, 7 dotted light blue). Decreases in dual task cost indicate performance improvement. **(B)** Averages across subjects of dual task cost values before and after Dance for PD, as well as before and after matched-intensity exercise. Mean values are reported, error bars indicate standard error (also in parenthesis). In red, the means across subjects of the differences between before and after for both Dance for PD and matched-intensity exercise, with standard error in parenthesis.

## 4 DISCUSSION

This study investigated the hypothesis that dance, compared to an exercise intervention of matched intensity, may yield different outcomes based on the artistic context of the movement experience using the same patients as their own control subjects in the two different interventions. In particular, Dance for PD classes may produce physiological, affective, self-efficacy related, and motor changes that differ from a matched-intensity exercise session lacking artistic elements like the presence of music, the use of narrative and metaphorical language, and a social reality of grace and beauty, established and reinforced by dance teachers, live musicians, and a group of peers with PD addressed as “dancers.”

This study aimed to investigate how the artistic elements of a Dance for PD sessions may shape therapeutic changes. The presence of teachers was purposely excluded from the matched-intensity exercise intervention to remove from this condition the potential therapeutic effects mediated by the relationship with the participants, contributing to their identification as “dancers,” “dance students,” or more in general “capable movers.” The enthusiasm and motivation cultivated by dance teachers or instructors, and experienced by dance participants, is integral to the (social) reality of art partaking, thus constituting an important feature of the artistic context potentially conducive to therapeutic outcomes.

### 4.1 Participants

The subjects who participated in this study were at a stages I or II of the Hoehn & Yahr scale (Goetz et al., 2004), although included a broad range of differences in terms of duration and clinical manifestations of the disease (MDS-UPDRS score), age, and levodopa administration. Because of the range of clinical characteristics in this population, the study was designed for within-subjects pre-post comparisons, investigating whether within the same subject there could be observable differences in response to a Dance for PD class and a matched-intensity exercise session. This within-subjects design was the most appropriate strategy to control for the heterogeneity of clinical features among participants in this real-world community setting (Greenland et al., 2019). Although this study did not include age-matched controls, future studies may question whether the described effects of Dance for PD classes apply specifically to those with PD or may be extended to an aging population in general.

### 4.2 Heart Rate (HR) and Electrodermal Activity (EDA)

Exercise prescriptions are defined by the frequency, intensity, duration, and type of exercise (aerobic, resistance, flexibility and neuromuscular training) recommended to the patients (Ignaszewski et al., 2017; Garber and Deschenes, 2017). This study compared the effects of a one-time movement session (Dance for PD, matched-intensity exercise) of equal duration and type. Importantly, the percentage heart rate response (PHRR) values were not significantly different between Dance for PD and matched-intensity exercise, thus it is appropriate to consider these two conditions equally demanding in terms of cardiovascular effort (Ignaszewski et al., 2017). Exercise intensity did not change in the two interventions, with PHRR values below 30% which are considered “very light” exercise (Ignaszewski et al., 2017), and the specific structure of the session remained the same, from mobilizing upper and lower joints while sitting on a chair, then transitioning to standing and finally walking in space.

However, electrodermal activity revealed differences between the two conditions, in terms of changes in the average skin conductance level (SCL) and in the amplitude of spontaneous fluctuations (SF). Significantly higher SCL and SF amplitude were observed during Dance for PD than during matched-intensity exercise. Electrodermal activity depends on the sweat glands function mediated by the sympathetic nervous system (Bach et al., 2010; Boucsein, 2012), which takes part in the homeostatic process of thermoregulation, as well as it is modulated by psychological factors, like emotional arousal and autonomic stress responses (Boucsein, 2012; Critchley, 2002). Evidence of higher SCL and SF amplitude during Dance for PD than during matched-intensity exercise may indicate increased sympathetic activation during Dance for PD. Importantly, skin conductance, differently from heart rate, is modulated by sympathetic activity only. Also, since both conditions share the same (“very light”) activity level, based on the PHRR analysis, it is unlikely that the activity of sweat glands would be exercise-dependent, as both interventions may involve similar processes of thermoregulation. Furthermore, the significant correlation between the general positive affect values calculated after the Dance for PD sessions, reported through the PANAS-X questionnaire, and the average skin conductance levels measured during the class (Figure 4) suggests that these differences in electrodermal activity may be primarily attributed to emotional arousal. In particular, EDA reflects changes in skin resistance in response to the activation of the sympathetic nervous system, controlling the permeability of sweat glands associated with thermoregulation processes, emotional arousal, and stress responses (Cecchi et al., 2020). As we compared two conditions with matching physical activity levels, taking place in the same environment with centralized thermostat control (i.e., The Mark Morris Dance Center), at the same time of the day, a couple of days apart, within the same subjects, we may rule out that the main explanation for the observed EDA changes is thermoregulation. Since the evidence of higher SCL and SF amplitude during dance is accompanied by self-reported increases in general positive affect, we may interpret the EDA signal in light of affective changes, attributing these differences in skin conductance to emotional arousal. It is important to remember that participants experienced a Dance for PD class taught by professional dancers with the use metaphorical language, live music, and dance peers sharing a reality of art partaking. All these features were removed from the matched-intensity exercise session, during which participants engaged with instructional commands from a computer screen. The context of the movement experience was intentionally altered and, while not affecting participants’ physical activity levels, this manipulation may be responsible for the differences in emotional arousal suggested by EDA changes and paralleled by the self-reported affective scores. Because of the limited size of this sample, it is necessary to collect further data in future studies in order to confirm this correlation once in-person classes commence.

### 4.3 Affect, Self-Efficacy, Beauty

The impact of dance interventions on mood disturbances in PD, including depression, anxiety, and apathy, has been previously reported by McNeely et al. (2015). Also, a recent systematic review confirmed the effects of dance movement therapy in reducing depression in adults in general (Karkou et al., 2019). In this study’s sample, subjects reported minimal to mild depressive symptoms (Table 3), assessed through the Beck Depression Inventory-II (BDI-II) at baseline. Importantly, depression is associated with both a reduction in positive affect, a broad dimension including a range of pleasant emotions like joy, interest, enthusiasm, alertness and self-confidence, as well as an increase in negative affect, including fear, anxiety, irritability, loneliness, hostility and shame (Nutt et al., 2007). In this study, with only seven subjects filling out the self-reported questionnaires, the trend of an increase in general positive affect (GPA) after Dance for PD failed to reach significance in front of the average decrease in GPA after matched-intensity exercise (Figure 3). However, GPA after Dance for PD was positively correlated with skin conductance levels during dance, which was significantly higher than during matched-intensity exercise. Since self-reported questionnaires are typically administered to larger groups of subjects (Demetriou et al., 2015), future studies should compare affective changes following these two interventions in a larger sample to increase statistical power to examine this trend.

In Koch et al. (2016) participants’ well-being, measured by the “Heidelberg State Inventory” and including tension, anxiety, coping, positive affect, depressed affect, and vitality, was found to be significantly higher after an Argentine tango intervention in a group of 34 patients with PD. In this study, the researchers reported increased well-being, body self-efficacy, and experience of beauty after a single tango class for people with PD. However, the study did not include a control condition but only a pre-post assessment of the dance intervention. In our sample, we were able to compare a single Dance for PD class and a matched-intensity exercise session using the same people with PD, finding significant differences in both body self-efficacy (BSE) and beauty subscale (Figure 5) between these two conditions. In particular, the beauty subscale represents a sub-dimension (aesthetic experience) within the BSE questionnaire (Fuchs and Koch, 2014) comprising two items that state, “My movements are beautiful” and “I can move elegantly/with grace” (Koch et al., 2016). The concepts of beauty and grace are often intertwined with dance (Houston, 2015) and may constitute prior knowledge (Bless & Greifeneder, 2018) of dance as the art of movement. However, it is important to emphasize that Dance for PD participants directly experience themselves as more graceful and beautiful (Houston, 2015), significantly more so than after matched-intensity exercise, as evidenced by the beauty subscale self-report. Both Houston (2015) and Koch (2017) suggest that the experience of feeling beautiful is foundational to self-efficacy, the belief in one’s own abilities, identity, and dignity. Further, Lewthwaite and Wulf (2017) argue that attentional and motivational mechanisms, the latter including both positive affect and self-efficacy, are central to enhancing motor performance (Lewthwaite and Wulf, 2017). Feeling beautiful may contribute both to the beliefs of what the body is capable of (self-efficacy) and to the affective value associated with movement (positive affect) thus, ultimately, to motor performance.

As previously described, Dance for PD classes were taught by dance professionals and live musicians, while the matched-intensity exercise sessions were delivered through instructional commands on a computer screen. We chose to remove any form of social interaction from the latter condition since we argue that the sense of beauty and capability is socially reinforced (Bless & Greifeneder, 2018) by the presence of teaching artists, as well as of fellow “dancers,” affirming and validating ability, grace, and beauty in each other (Houston, 2015).

### 4.4 Gait Symmetry and Dual Task Performance

The role of dance practice in reducing PD motor symptoms has been previously reviewed (Carvalho Aguiar et al., 2016; Sharp and Hewitt, 2014) showing the beneficial effects of dance on motor symptoms (UPDRS 3), balance (Berg balance scale), and walking velocity (6-minute walking test). Hashimoto at al. (2015) reported effects on both motor (Berg balance scale, Timed-Up-and-Go test) and cognitive functions (Frontal Assessment Battery, Mental Rotation Task). Decline in motor performance and cognitive impairment have the greatest influence on the quality of life of people with PD (Shrag et al., 2000).

In our sample, two subjects reported considerable issues with gait, falls and freezing, while two subjects scored between 24 and 25 (cutoff point) on the Montreal Cognitive Assessment (MoCA) (Table 3). Gait performance over the 6-minute walking test was analyzed through a portable system for functional assessment of movement (GWALK). The symmetry index (Figure 6), providing a comparison of the anteroposterior acceleration between the right and left gait cycles, revealed significant differences between the Dance for PD and matched-intensity exercise interventions. After dance, participants showed a significantly higher symmetry of gait than after exercise. Importantly, Yogev et al. (2007) reported that, differently from healthy older adults, PD patients rely on attention and cognitive resources to maintain a bilaterally coordinated gait, and individuals who are more prone to falling score lower on executive function tasks and attention indexes. Importantly, gait symmetry has been related to freezing of gait and fear of falling (Frazzitta et al., 2013), which is an essential factor in determining balance, posture, and functional mobility (Franchignoni et al., 2015). Reduced attention has been linked to increased fall frequency in PD (Allcock et al., 2008). The observed increase in gait symmetry after Dance for PD, compared to matched-intensity exercise, may be explained by a increased executive function or an increased attention to walking. This finding is important in light of the significance of gait symmetry in the control of balance and its relationship to fall risk.

When testing patients during simultaneous motor and cognitive tasks, Rochester et al. (2004) reported a decrease in gait velocity compared to the single (motor) task condition in subjects with PD more than controls. The study interpreted this evidence as a result of the competition for attentional resources, particularly when cognitive resources were limited, as in patients with PD presenting executive dysfunction or mild cognitive impairment. Dual task performance can be achieved by constantly shifting or dividing attention between two tasks (Rochester et al., 2004). Difficulty maintaining the motor performance may be due to the allocation of attentional resources toward the cognitive task, thus decreasing one’s attention to gait. Subjects with PD have increased demands to cognitively attend to movements like walking that were previously considered automatic (Rochester et al., 2004). Importantly, the current study reports an improvement in gait performance during a cognitive task (reduction in dual task cost) significantly higher after Dance for PD than after matched-intensity exercise (Figure 7). In line with this finding, a recent review on the effects of dance in PD (Kalyani et al., 2019) highlighted the improvements in gait velocity during cognitive dual tasks following dance interventions, pointing out the lack of controls among the existing studies. The present research constitutes the first evidence that the enhanced dual task performance after Dance for PD was significantly different than after matched-intensity exercise (Figure 7) in a sample of subjects that completed both conditions (Dance for PD and matched-intensity exercise). The decrease in dual task cost may be explained with more efficient divided attention processes. However, it is also possible that after dance (but not after matched-intensity exercise) the need to cognitively attend to walking was reduced by increased gait automaticity, so that attentional resources could be allocated to the cognitive task with a smaller impact on gait performance. Automaticity could have been enhanced by an external focus of attention, an idea known as the “constrained action hypothesis” (Wulf et al., 2001; Kal et al., 2013), employed for the reduction of cognitive interference in athletic performance. Maintaining an external focus of attention, like a target or an image, has been shown to be an effective strategy in improving postural control in patients with PD (Wulf et al., 2009; Jazaeri et al., 2018), however no data has been reported on the effects of an external focus on dual task or gait performance. In dance practice, the use of metaphors may serve the purpose of directing attention externally to produce a movement through the use of imagery (Guss-West and Wulf, 2016; Lewthwaite and Wulf, 2017). Whether such images could provide a source of external focus to patients with PD, enhancing movement automaticity, needs to be further investigated. Future studies should explore the relationship between gait automaticity and external attentional focus in PD.

Further, gait initiation can be modulated by the presentation of emotional (pleasant or unpleasant) visual stimuli in patients with or without freezing of gait (FOG) (Lagravinese et al., 2018), as evidenced by longer reaction times and shorter steps in response to incongruent/unpleasant condition (stepping forward toward an unpleasant stimulus). This evidence may suggest an interplay between gait performance and emotional processing, since the basal ganglia act as gatekeepers in both approach motivation (Ikemoto et al., 2015) and avoidance responses (Hormigo et al., 2016). This interplay may be relevant to this study, as we reported affective responses paralleled by electrodermal activity, which significantly differed between the two conditions. In particular, after Dance for PD, participants showed higher emotional arousal (EDA and positive affect), increased gait symmetry, and faster motor performance during a dual task condition.

Importantly, the Timed-Up-and-Go test with the additional cognitive task is considered a valid tool to identify fall risk in people with PD (Vance et al., 2015), thus, the present finding has important implications on the fall risk, mobility and autonomy of the participants overtime.

### 4.5 Conclusion

This study explores the hypothesis that dance, compared to an exercise intervention of matched intensity, may yield different physiological, affective, self-efficacy, and motor effects. Using the same patients as their own controls, we compared a Dance for PD class with a matched-intensity exercise session designed ad-hoc to be lacking dance elements like music, metaphorical language, and social reality of grace and beauty shared with dance teachers and peers. We showed that dance yields different outcomes suggesting a possible interaction between affective responses, the experience of beauty, self-efficacy, and gait performance, traditionally separated in non-motor and motor features. The specific relationship to movement elicited by dancing may be responsible for the modulation of motivational and attentional mechanisms that influence motor behavior and can result in symptoms’ improvement in people with PD. In Dance for PD, thoughts involving distress and worry, may be at times “blissfully absent” (Gail Montero, 2016), distancing the person with PD from a perception of disability while reclaiming a sense of efficacy, personhood, and agency (Houston, 2015). We argue this process of perspective-transformation within the artistic context of a dance class not only to be relevant to psychological wellbeing (affect, self-efficacy), but also to have important functional implications. Improvement in the motor performance showed by patients with PD after dance practice (gait symmetry, dual task performance) may stem from affective modulation and decreased cognitive interference (diagnosis-related self-talk), potentially resulting in increased availability of attentional resources toward motor performance (Ferrazzoli et al., 2018, 2020), or in increased external focus enhancing movement automatization when needed (Kal et al., 2013). Future studies may explore these potential mechanisms to shed light on the specific processes that make dance a valuable resource for people with PD.

### 4.6 Study Limitations

Limitations of this study must be acknowledged. Although we employed a within-subjects design, this research involved a very limited sample of subjects with PD, so the ability to generalize to the population as a whole is limited. Because of the small size of this sample, it will be necessary to collect further data in future studies in order to confirm the reported findings. Further, this self-selected group regularly takes part in Dance for PD classes and willingly volunteered to participate in this study. It was not possible to recruit participants naïve to dance because of lack of volunteers’ availability. Participants were familiar with the dance intervention but not with the matched-intensity exercise condition, which might have introduced a bias toward dance in the analysis. Since the study took place in a “real world” community setting, it was not possible to fully balance the order of the two conditions (Dance for PD and matched-intensity exercise) due to the participants’ schedule. Similarly, we could not apply a wash out period between the two conditions, since it would have meant to remove the participants from the community that they were part of as active members. There was no way to conceal whether dance or exercise were being presented, therefore neither the subjects nor the researcher were blind to the type of intervention; however, participants were not aware of the research questions investigated in the study. Future studies should not only expand the sample size to confirm these results and allow a broader generalization but should also include dance naïve participants and explore the qualitative differences between the two modalities presented (Dance for PD and matched-intensity exercise). A mixed-methods research design, including a larger sample, appropriate randomization, and both dance and exercise naïve participants should follow our study to further investigate the hypotheses and results here reported.

## 8 Author Contributions

Fontanesi, C., planned and designed the study, collected the data, performed the analysis, interpreted data, and drafted the manuscript. DeSouza, J.F.X., contributed analysis tools, edited the draft, and approved the version to be submitted to the journal.

## 10 Acknowledgments

The authors thank David Leventhal, Maria Portman Kelly, The Mark Morris Dance Center, and the participants who volunteered to take part in this study. This would have not been possible without their generous collaboration and support. We thank Massimiliano Gobbo and Laura Vacchi at University of Brescia (Italy), as well as Timothy Ellmore and the research assistants at The City College of New York (CUNY) for their invaluable help in developing this study.

## 11 Data Availability Statement

The datasets generated for this study are available on request to the corresponding author.

## 12 Ethics Statement

The design and methodology of this study were approved by The City College of New York Institutional Review Board (IRB).

## References

Allcock, L. M., Rowan, E. N., Steen, I. N., Wesnes, K., Kenny, R. A., & Burn, D. J. (2009). Impaired attention predicts falling in Parkinson’s disease. Parkinsonism & related disorders, 15(2), 110–115. doi: 10.1016/j.parkreldis.2008.03.010

Bach, D.R., Friston, K.J., & Dolan R.J. (2010). Analytic Measures for Quantification of Arousal from Spontaneous Skin Conductance Fluctuations. International Journal of Psychophysiology, 76(1), 52–55.

Barnstaple, R., Hackney, M.E., Fontanesi, C., & DeSouza, J.F.X. (2018). Mechanisms of Dance in the Rehabilitation of Neurodegenerative Conditions. Brain, Body, Cognition 2018; 8(1):17–28.

Bautmans, I., Lambert, M., & Mets, T. (2004). The six-minute walk test in community dwelling elderly: influence of health status. BMC geriatrics, 4, 6. doi:10.1186/1471-2318-4-6

Bearss, K., McDonald, K.C., Bar, R.J. & DeSouza, J.F.X. (2017). A 12 week Dance Intervention for Parkinson’s Disease: motor functions and quality of life. Advances in Integrative Medicine. doi:10.1016/j.aimed.2017.02.002

Beck, A.T., Steer, R.A., & Brown, G.K. (1996). Manual for the Beck Depression Inventory-II. San Antonio, Texas: Psychol. Corp.

Bezner, J., & Rose, L. (1989). Adult exercise instruction sheets: Home exercises for rehabilitation. San Antonio, Texas: Therapy Skill Builders.

Bless, H., & Greifeneder, R. (2018). General framework of social cognitive processing. In Greifeneder, Bless. Social Cognition: How Individuals Construct Social Reality. 2nd ed. London: Routledge.

Boucsein, W. (2012). “Chapter 1: Principles of Electrodermal Phenomena,” in Electrodermal Activity (2nd ed.). (New York, NY: Springer US), 1–86. doi:10.1007/978-1-4614-1126-0

Braithwaite, J.J., Watson, D.G., Jones, R., & Rowe M. (2015). A Guide for Analysing Electrodermal Activity (EDA) & Skin Conductance Responses (SCRs) for Psychological Experiments. https://www.birmingham.ac.uk/Documents/college-les/psych/saal/guide-electrodermal-activity.pdf [Accessed August 15, 2019].

Bryant, M. S., Workman, C. D., Hou, J. G., Henson, H. K., & York, M. K. (2016). Acute and Long-Term Effects of Multidirectional Treadmill Training on Gait and Balance in Parkinson Disease. PM & R: the journal of injury, function, and rehabilitation, 8(12), 1151–1158. https://doi.org/10.1016/j.pmrj.2016.05.001

BTS Bioengineering (2016). G-WALK User Manual English Version 8.0.0. Brooklyn, NY: BTS Bioengineering Corp.

Carvalho Aguiar, L.P., Alves da Rocha, P., Morris, M. (2016). Therapeutic Dancing for Parkinson’s Disease. International Journal of Gerontology, 10(2), 64–70, doi:10.1016/j.ijge.2016.02.002.

Cecchi, S., Piersanti, A., Poli, A. and Spinsante, S. (2020). Physical Stimuli and Emotions: EDA Features Analysis from a Wrist-Worn Measurement Sensor. IEEE 25th International Workshop on Computer Aided Modeling and Design of Communication Links and Networks (CAMAD), 1–6, doi: 10.1109/CAMAD50429.2020.9209307.

Chou, K.L., Lenhart, A., Koeppe, R.A., & Bohnen N.I. (2014). Abnormal MoCA and Normal Range MMSE Scores in Parkinson Disease without Dementia: Cognitive and Neurochemical Correlates. Parkinsonism & Related Disorders, 20(10), 1076–80. doi:10.1016/j.parkreldis.2014.07.008

Ciantar, S.R., Bearss, K.A., Levkov, G., Bar, R.J., DeSouza, J.F.X. (2019). Investigating Affective and Motor Improvements with Dance in Parkinson’s Disease. BioRxiv 665711; doi: https://doi.org/10.1101/665711

Critchley, H. D. (2002). Review: Electrodermal Responses: What Happens in the Brain. The Neuroscientist, 8(2), 132–142. doi:10.1177/107385840200800209

de Natale, E. R., Paulus, K. S., Aiello, E., Sanna, B., Manca, A., Sotgiu, G., Leali, P. T., & Deriu, F. (2017). Dance therapy improves motor and cognitive functions in patients with Parkinson’s disease. NeuroRehabilitation, 40(1), 141–144. doi:10.3233/NRE-161399

de Oliveira Segundo, V.H., Brandao de Albuquerque Filho, N.J. Mendes Rebouças, G., Renee Felipe, T., Ferreira Matos, V.R., Silva Dantas, P.M., Fonseca Pinto, E. (2016). Use of predictive equations of maximum heart rate for exercise prescription: a comparative study. Journal of Sports and Physical Education (IOSR-JSPE), 3(1), 04–08. doi:10.9790/6737-0310408

Demetriou, C., Özer, B., Essau, C. (2015). “Self-Report Questionnaires,” in The Encyclopedia of Clinical Psychology, First Edition, ed. R.L. Cautin and S.O. Lilienfeld (John Wiley & Sons, Inc.). doi: 10.1002/9781118625392.wbecp507

DeSouza, J.F.X., Bearss, K. (2018). Progression of Parkinson’s disease symptoms halted using dance over 3-years as assessed with MDS-UPDRS. Neuroscience Meeting Planner. San Diego: Society for Neuroscience Abstracts. doi: https://abstractsonline.com/pp8/#!/4649/presentation/22209

Duncan, R. P., & Earhart, G. M. (2012). Randomized controlled trial of community-based dancing to modify disease progression in Parkinson disease. Neurorehabilitation and neural repair, 26(2), 132–143. doi:10.1177/1545968311421614

Earhart, G. M., Duncan, R. P., Huang, J. L., Perlmutter, J. S., & Pickett, K. A. (2015). Comparing interventions and exploring neural mechanisms of exercise in Parkinson disease: a study protocol for a randomized controlled trial. BMC neurology, 15, 9. doi:10.1186/s12883-015-0261-0

Empatica. (2018). E4 Wristband from Empatica user’s manual #E4-SP069-B-20150001. https://empatica.app.box.com/v/E4-User-Manual [Accessed August 17, 2018].

Ferrazzoli, D., Ortelli, P., Cucca, A., Bakdounes, L., Canesi, M., & Volpe, D. (2020). Motor-cognitive approach and aerobic training: a synergism for rehabilitative intervention in Parkinson’s disease. Neurodegenerative disease management, 10(1), 41–55. doi:10.2217/nmt-2019-0025

Ferrazzoli, D., Ortelli, P., Madeo, G., Giladi, N., Petzinger, G. M., & Frazzitta, G. (2018). Basal ganglia and beyond: The interplay between motor and cognitive aspects in Parkinson’s disease rehabilitation. Neuroscience and biobehavioral reviews, 90, 294–308. doi:10.1016/j.neubiorev.2018.05.007

Foster, E. R., Golden, L., Duncan, R. P., & Earhart, G. M. (2013). Community-based Argentine tango dance program is associated with increased activity participation among individuals with Parkinson’s disease. Archives of physical medicine and rehabilitation, 94(2), 240–249. doi:10.1016/j.apmr.2012.07.028

Franchignoni, F., Martignoni, E., Ferriero, G., & Pasetti, C. (2005). Balance and fear of falling in Parkinson’s disease. Parkinsonism & related disorders, 11(7), 427–433. doi:10.1016/j.parkreldis.2005.05.005

Frazzitta, G., Balbi, P., Maestri, R., Bertotti, G., Boveri, N., & Pezzoli, G. (2013). The beneficial role of intensive exercise on Parkinson disease progression. American journal of physical medicine & rehabilitation, 92(6), 523–532. doi:10.1097/PHM.0b013e31828cd254

Frazzitta, G., Maestri, R., Bertotti, G., Riboldazzi, G., Boveri, N., Perini, M., Uccellini, D., Turla, M., Comi, C., Pezzoli, G., & Ghilardi, M. F. (2015). Intensive rehabilitation treatment in early Parkinson’s disease: a randomized pilot study with a 2-year follow-up. Neurorehabilitation and neural repair, 29(2), 123–131. https://doi.org/10.1177/1545968314542981

Frazzitta, G., Pezzoli, G., Bertotti, G., & Maestri, R. (2013). Asymmetry and freezing of gait in parkinsonian patients. Journal of neurology, 260(1), 71–76. doi:10.1007/s00415-012-6585-4

Fuchs, T., & Koch, S. C. (2014). Embodied affectivity: on moving and being moved. Frontiers in psychology, 5, 508. doi:10.3389/fpsyg.2014.00508

Gail Montero, B. (2016). The Pleasure of Movement and the Awareness of the Self. In Gail Montero, B. (Ed.), Thought in Action: Expertise and the Conscious Mind. Oxford Scholarship Online. doi:10.1093/acprof:oso/9780199596775.001.0001

Garbarino, M., Lai, M., Bender, D., Picard R.W. & Tognetti, S. (2014). Empatica E3 — A wearable wireless multi-sensor device for real-time computerized biofeedback and data acquisition. 2014 4th International Conference on Wireless Mobile Communication and Healthcare-Transforming Healthcare Through Innovations in Mobile and Wireless Technologies (MOBIHEALTH), Athens. 39–42, doi:10.1109/MOBIHEALTH.2014.7015904

Garber, C.E., Deschenes, M.D. (2017). “Chapter 5: General Principles of Exercise Prescription,” in ACSM’s Guidelines for Exercise Testing and Prescription, ed. American College of Sports Medicine (Philadelphia, PA: Lippincott, Williams & Wilkins).

Giladi, N., Shabtai, H., Simon, E. S., Biran, S., Tal, J., & Korczyn, A. D. (2000). Construction of freezing of gait questionnaire for patients with Parkinsonism. Parkinsonism & Related Disorders, 6(3), 165–170. PubMed PMID: 10817956

Greenland, J.C., Williams-Gray, C.H., Barker, R.A. (2019) The clinical heterogeneity of Parkinson’s disease and its therapeutic implications. Eur J Neurosci, 49(3):328–338. doi: 10.1111/ejn.14094. Epub 2018 Oct 14. PMID: 30059179.

Guss-West, C., & Wulf, G. (2016). Attentional Focus in Classical Ballet: A Survey Of Professional Dancers. Journal of dance medicine & science: official publication of the International Association for Dance Medicine & Science, 20(1), 23–29. doi:10.12678/1089-313X.20.1.23

Hackney, M. E., & Earhart, G. M. (2009). Effects of dance on movement control in Parkinson’s disease: a comparison of Argentine tango and American ballroom. Journal of rehabilitation medicine, 41(6), 475–481. doi:10.2340/16501977-0362

Hackney, M. E., & Earhart, G. M. (2010). Effects of dance on gait and balance in Parkinson’s disease: a comparison of partnered and nonpartnered dance movement. Neurorehabilitation and neural repair, 24(4), 384–392. doi:10.1177/1545968309353329

Hackney, M. E., Kantorovich, S., Levin, R., & Earhart, G. M. (2007). Effects of tango on functional mobility in Parkinson’s disease: a preliminary study. Journal of neurologic physical therapy: JNPT, 31(4), 173–179. doi:10.1097/NPT.0b013e31815ce78b

Hashimoto, H., Takabatake, S., Miyaguchi, H., Nakanishi, H., & Naitou, Y. (2015). Effects of dance on motor functions, cognitive functions, and mental symptoms of Parkinson’s disease: a quasi-randomized pilot trial. Complementary therapies in medicine, 23(2), 210–219. doi:10.1016/j.ctim.2015.01.010

Heiberger, L., Maurer, C., Amtage, F., Mendez-Balbuena, I., Schulte-Mönting, J., Hepp-Reymond, M. C., & Kristeva, R. (2011). Impact of a weekly dance class on the functional mobility and on the quality of life of individuals with Parkinson’s disease. Frontiers in aging neuroscience, 3, 14. doi:10.3389/fnagi.2011.00014

Hymmen, P., Stalker, C. A., & Cait, C. A. (2013). The case for single-session therapy: does the empirical evidence support the increased prevalence of this service delivery model?. Journal of mental health (Abingdon, England), 22(1), 60–71. https://doi.org/10.3109/09638237.2012.670880

Hoops, S., Nazem, S., Siderowf, A.D., Duda, J.E., Xie, S.X., Stern, M.B., & Weintraub, D. (2009). Validity of the MoCA and MMSE in the Detection of MCI and Dementia in Parkinson Disease. Neurology, 73, 1738–1745. doi: 10.1212/WNL.0b013e3181c34b47

Hormigo, S., Vega-Flores, G., and Castro-Alamancos, M. A. (2016). Basal Ganglia Output Controls Active Avoidance Behavior. The Journal of neuroscience: the official journal of the Society for Neuroscience, 36(40), 10274–10284. doi:10.1523/JNEUROSCI.1842-16.2016

Houston, S. (2015). Feeling Lovely: An Examination of the Value of Beauty for People Dancing with Parkinson’s. Dance Research Journal, 47(1), 27–43. doi:10.1017/S0149767715000042

Houston, S., & McGill, A. (2013). A mixed-methods study into ballet for people living with Parkinson’s. Arts & health, 5(2), 103–119. doi:10.1080/17533015.2012.745580

Ignaszewski, M., Lau, B., Wong, S., Isserow, S. (2017). The science of exercise prescription: Martti Karvonen and his contributions. British Columbia Medical Journal (BCMJ), 59(1):38–41.

Ikemoto, S., Yang, C., and Tan, A. (2015). Basal ganglia circuit loops, dopamine and motivation: A review and enquiry. Behavioural brain research, 290, 17–31. doi:10.1016/j.bbr.2015.04.018

Jazaeri, S. Z., Azad, A., Mehdizadeh, H., Habibi, S. A., Mandehgary Najafabadi, M., Saberi, Z. S., Rahimzadegan, H., Moradi, S., Behzadipour, S., Parnianpour, M., Taghizadeh, G., & Khalaf, K. (2018). The effects of anxiety and external attentional focus on postural control in patients with Parkinson’s disease. PloS one, 13(2), e0192168. doi:10.1371/journal.pone.0192168

Kal, E. C., van der Kamp, J., & Houdijk, H. (2013). External attentional focus enhances movement automatization: a comprehensive test of the constrained action hypothesis. Human movement science, 32(4), 527–539. doi:10.1016/j.humov.2013.04.001

Kalyani, H., Sullivan, K., Moyle, G., Brauer, S., Jeffrey, E. R., Roeder, L., Berndt, S., & Kerr, G. (2019). Effects of Dance on Gait, Cognition, and Dual-Tasking in Parkinson’s Disease: A Systematic Review and Meta-Analysis. Journal of Parkinson’s disease, 9(2), 335–349. doi:10.3233/JPD-181516

Karkou, V., Aithal, S., Zubala, A., & Meekums, B. (2019). Effectiveness of Dance Movement Therapy in the Treatment of Adults With Depression: A Systematic Review With Meta-Analyses. Frontiers in psychology, 10, 936. doi:10.3389/fpsyg.2019.00936

Koch, S. C., Mergheim, K., Raeke, J., Machado, C. B., Riegner, E., Nolden, J., Diermayr, G., von Moreau, D., & Hillecke, T. K. (2016). The Embodied Self in Parkinson’s Disease: Feasibility of a Single Tango Intervention for Assessing Changes in Psychological Health Outcomes and Aesthetic Experience. Frontiers in neuroscience, 10, 287. doi:10.3389/fnins.2016.00287

Koch, S.C. (2017). Arts and health: Active factors and a theory framework of embodied aesthetics. The Arts in Psychotherapy, 54, 85–91. doi:10.1016/j.aip.2017.02.002

Kunkel, D., Fitton, C., Roberts, L., Pickering, R. M., Roberts, H. C., Wiles, R., Hulbert, S., Robison, J., & Ashburn, A. (2017). A randomized controlled feasibility trial exploring partnered ballroom dancing for people with Parkinson’s disease. Clinical rehabilitation, 31(10), 1340–1350. doi:10.1177/0269215517694930

Lagravinese, G., Pelosin, E., Bonassi, G., Carbone, F., Abbruzzese, G., and Avanzino, L. (2018). Gait initiation is influenced by emotion processing in Parkinson’s disease patients with freezing. Movement disorders: official journal of the Movement Disorder Society, 33(4), 609–617. doi:10.1002/mds.27312

Lewthwaite, R., & Wulf, G. (2017). Optimizing motivation and attention for motor performance and learning. Current opinion in psychology, 16, 38–42. doi:10.1016/j.copsyc.2017.04.005

McGill, A., Houston, S., & Lee, R. Y. (2014). Dance for Parkinson’s: a new framework for research on its physical, mental, emotional, and social benefits. Complementary therapies in medicine, 22(3), 426–432. doi:10.1016/j.ctim.2014.03.005

McKee, K. E., & Hackney, M. E. (2013). The effects of adapted tango on spatial cognition and disease severity in Parkinson’s disease. Journal of motor behavior, 45(6), 519–529. doi:10.1080/00222895.2013.834288

McNeely, M. E., Duncan, R. P., & Earhart, G. M. (2015). Impacts of dance on non-motor symptoms, participation, and quality of life in Parkinson disease and healthy older adults. Maturitas, 82(4), 336–341. doi:10.1016/j.maturitas.2015.08.002

Müller, P., Rehfeld, K., Schmicker, M., Hökelmann, A., Dordevic, M., Lessmann, V., Brigadski, T., Kaufmann, J., & Müller, N. G. (2017). Evolution of Neuroplasticity in Response to Physical Activity in Old Age: The Case for Dancing. Frontiers in aging neuroscience, 9, 56. https://doi.org/10.3389/fnagi.2017.00056

Nutt, D., Demyttenaere, K., Janka, Z., Aarre, T., Bourin, M., Canonico, P. L., Carrasco, J. L., & Stahl, S. (2007). The other face of depression, reduced positive affect: the role of catecholamines in causation and cure. Journal of psychopharmacology (Oxford, England), 21(5), 461–471. doi:10.1177/0269881106069938

Oberfeld, D., Franke, T. Evaluating the robustness of repeated measures analyses: The case of small sample sizes and nonnormal data. Behav Res 45, 792–812 (2013). doi:10.3758/s13428-012-0281-2

Peper, E., Shaffer, F., & Lin, I.M. (2010). Garbage In; Garbage Out—Identify Blood Volume Pulse (BVP) Artifacts Before Analyzing and Interpreting BVP, Blood Volume Pulse Amplitude, and Heart Rate/Respiratory Sinus Arrhythmia Data. Biofeedback, 38(1), 19–23. doi:10.5298/1081-5937-38.1.19

Petzinger, G. M., Fisher, B. E., McEwen, S., Beeler, J. A., Walsh, J. P., & Jakowec, M. W. (2013). Exercise-enhanced neuroplasticity targeting motor and cognitive circuitry in Parkinson’s disease. The Lancet. Neurology, 12(7), 716–726. doi:10.1016/S1474-4422(13)70123-6

Podsiadlo, D., & Richardson, S. (1991). The timed ‘Up & Go’: A test of basic functional mobility for frail elderly persons. Journal of the American Geriatrics Society, 39(2), 142–148. doi: 10.1111/j.1532-5415.1991.tb01616.x

Rehfeld, K., Lüders, A., Hökelmann, A., Lessmann, V., Kaufmann, J., Brigadski, T., Müller, P., & Müller, N. G. (2018). Dance training is superior to repetitive physical exercise in inducing brain plasticity in the elderly. PloS one, 13(7), e0196636. https://doi.org/10.1371/journal.pone.0196636

Rehfeld, K., Müller, P., Aye, N., Schmicker, M., Dordevic, M., Kaufmann, J., Hökelmann, A., & Müller, N. G. (2017). Dancing or Fitness Sport? The Effects of Two Training Programs on Hippocampal Plasticity and Balance Abilities in Healthy Seniors. Frontiers in human neuroscience, 11, 305. https://doi.org/10.3389/fnhum.2017.00305

Rocha, P. A., Porfírio, G. M., Ferraz, H. B., & Trevisani, V. F. (2014). Effects of external cues on gait parameters of Parkinson’s disease patients: a systematic review. Clinical neurology and neurosurgery, 124, 127–134. doi:10.1016/j.clineuro.2014.06.026

Rochester, L., Hetherington, V., Jones, D., Nieuwboer, A., Willems, A. M., Kwakkel, G., & Van Wegen, E. (2004). Attending to the task: interference effects of functional tasks on walking in Parkinson’s disease and the roles of cognition, depression, fatigue, and balance. Archives of physical medicine and rehabilitation, 85(10), 1578–1585. doi:10.1016/j.apmr.2004.01.025

Schenkman, M., Moore, C. G., Kohrt, W. M., Hall, D. A., Delitto, A., Comella, C. L., Josbeno, D. A., Christiansen, C. L., Berman, B. D., Kluger, B. M., Melanson, E. L., Jain, S., Robichaud, J. A., Poon, C., & Corcos, D. M. (2018). Effect of High-Intensity Treadmill Exercise on Motor Symptoms in Patients With De Novo Parkinson Disease: A Phase 2 Randomized Clinical Trial. JAMA Neurol. 75(2):219–226. doi:10.1001/jamaneurol.2017.3517

Schrag, A., Jahanshahi, M., & Quinn, N. (2000). What contributes to quality of life in patients with Parkinson’s disease?. Journal of neurology, neurosurgery, and psychiatry, 69(3), 308–312. doi:10.1136/jnnp.69.3.308

Sharp, K., & Hewitt, J. (2014). Dance as an intervention for people with Parkinson’s disease: a systematic review and meta-analysis. Neuroscience and biobehavioral reviews, 47, 445–456. doi:10.1016/j.neubiorev.2014.09.009

Shumway-Cook, A., Brauer, S., & Woollacott, M. (2000). Predicting the Probability for Falls in Community-Dwelling Older Adults Using the Timed Up & Go Test. Physical Therapy, 80(9), 896–903.

Soleymani, M., Villaro-Dixon, F., Pun, T., & Chanel, G. (2017). Toolbox for Emotional Feature Extraction from Physiological Signals (TEAP). Frontiers in ICT, 4:1. doi:10.3389/fict.2017.00001

Tomlinson, C.L., Stowe, R., Patel, S., Rick, C., Gray, R., & Clarke, C.E. (2010). Systematic Review of Levodopa Dose Equivalency Reporting in Parkinson’s Disease: Systematic Review of LED Reporting in PD. Movement Disorders, 25(15), 2649–53. doi:10.1002/mds.23429

UK National Health Service. (2018). Physical activity guidelines for older adults-Exercise. https://www.nhs.uk/live-well/exercise/physical-activity-guidelines-older-adults/ [Accessed January 16, 2018]

Vance, R. C., Healy, D. G., Galvin, R., & French, H. P. (2015). Dual tasking with the timed “up & go” test improves detection of risk of falls in people with Parkinson disease. Physical therapy, 95(1), 95–102. doi:10.2522/ptj.20130386

Volpe, D., Signorini, M., Marchetto, A., Lynch, T., & Morris, M. E. (2013). A comparison of Irish set dancing and exercises for people with Parkinson’s disease: a phase II feasibility study. BMC geriatrics, 13, 54. doi:10.1186/1471-2318-13-54

Watson, D., & Clark, L. (1994). The PANAS-X: Manual for the Positive and Negative Affect Schedule-Expanded Form. University of Iowa. doi:10.17077/48vt-m4t2

Westheimer, O., McRae, C., Henchcliffe, C., Fesharaki, A., Glazman, S., Ene, H., & Bodis-Wollner, I. (2015). Dance for PD: a preliminary investigation of effects on motor function and quality of life among persons with Parkinson’s disease (PD). Journal of neural transmission (Vienna, Austria: 1996), 122(9), 1263–1270. doi:10.1007/s00702-015-1380-x

Wulf, G., Landers, M., Lewthwaite, R., & Töllner, T. (2009). External focus instructions reduce postural instability in individuals with Parkinson disease. Physical therapy, 89(2), 162–168. doi:10.2522/ptj.20080045

Wulf, G., McNevin, N., & Shea, C. H. (2001). The automaticity of complex motor skill learning as a function of attentional focus. The Quarterly journal of experimental psychology. A, Human experimental psychology, 54(4), 1143–1154. doi:10.1080/713756012

Yogev, G., Plotnik, M., Peretz, C., Giladi, N., & Hausdorff, J. M. (2007). Gait asymmetry in patients with Parkinson’s disease and elderly fallers: when does the bilateral coordination of gait require attention?. Experimental brain research, 177(3), 336–346. doi:10.1007/s00221-006-0676-3

