## Supplementary material for "Beauty That Moves: Dance for Parkinson’s Effects on Affect, Self-Efficacy, Gait Symmetry and Dual Task Performance": Table S1

### 1 Supplementary Figures and Tables

| | Sit to Stand (s) | | Delay before Forward Gait (s) | | Forward Gait (s) | |
|---|---|---|---|---|---|---|
| Subject ID | DfPD | MIE | DfPD | MIE | DfPD | MIE |
| 2 | -0.700 | -0.280 | -0.020 | 2.010 | 0.060 | -3.220 |
| 3 | -0.245 | -0.300 | 0.040 | 1.040 | -0.245 | -0.210 |
| 4 | -0.200 | -0.050 | 0.140 | 0.080 | 0.080 | -0.410 |
| 5 | -0.005 | 0.160 | -0.330 | 0.050 | -0.325 | 0.140 |
| 7 | -0.120 | 0.205 | 0.060 | 0.520 | 0.165 | -1.330 |
| **mean** | **-0.254** | **-0.053** | **-0.022** | **0.740** | **-0.053** | **-1.006** |
| **SD** | **0.265** | **0.237** | **0.181** | **0.816** | **0.217** | **1.352** |
| | Mid Turning (s) | | Return Gait (s) | | End Turning & Stand to Sit (s) | |
| Subject ID | DfPD | MIE | DfPD | MIE | DfPD | MIE |
| 2 | -0.520 | 1.320 | -2.010 | -1.330 | 0.060 | 0.630 |
| 3 | -0.275 | 0.770 | -0.595 | 0.555 | -2.475 | -0.380 |
| 4 | -0.450 | -0.380 | -0.675 | 0.660 | 0.555 | -0.185 |
| 5 | -0.030 | -0.535 | 0.005 | -0.405 | 0.065 | 0.350 |
| 7 | -0.905 | 0.345 | 0.350 | -1.035 | -0.860 | 0.430 |
| **mean** | **-0.436** | **0.304** | **-0.585** | **-0.311** | **-0.531** | **0.169** |
| **SD** | **0.323** | **0.778** | **0.903** | **0.903** | **1.201** | **0.430** |

**Table S1**: Changes in dual task cost (After-Before) intervention, in both Dance for PD (DfPD) and matched-intensity exercise (MIE). Across subjects means and standard deviations (SD) in bold.
